# Sorting nexin 5 mediates antigen presentation and immunity against *Mycobacterium tuberculosis*

**DOI:** 10.64898/2026.01.30.702915

**Authors:** Beatriz R. S. Dias, Kubra F. Naqvi, Victoria A. Ektnitphong, Samuel Alvarez-Arguedas, Kathryn C. Rahlwes, Priscila C. Campos, Michael U. Shiloh

**Author notes:** Corresponding author: Michael U. Shiloh, MD, PhD, Department of Medicine, Division of Infectious Diseases Department of Microbiology University of Texas Southwestern 5323 Harry Hines Blvd., Y9.308 Dallas, TX 75390-9113.

## Abstract

Tuberculosis (TB) remains one of the leading causes of death from a single infectious agent worldwide, yet the host pathways that regulate antigen presentation and lung inflammation during *Mycobacterium tuberculosis* (Mtb) infection are incompletely defined. Sorting nexin 5 (SNX5) is a protein best known for roles in endosomal trafficking, antigen processing, and antiviral host defense, but its contribution to immunity during Mtb infection is unknown. Here, we show that SNX5-deficient mice exhibit markedly increased mortality following low-dose aerosol infection despite unchanged pulmonary bacterial burden compared to wild-type mice. *Snx5*^−/−^ mice developed exacerbated lung inflammation without major alterations in immune cell recruitment. In macrophages, *Snx5* did not affect phagocytosis, vacuolar maturation, intracellular bacterial control, or global transcriptional responses to Mtb, but was required for efficient MHC class II antigen presentation. *Snx5* deficiency reduced antigen degradation, limited peptide loading onto MHC-II and impaired activation of antigen-specific CD4^+^ T cells without altering surface MHC-II abundance or expression of costimulatory molecules. Together, these findings identify SNX5 as a previously unrecognized regulator of MHC-II peptide loading that shapes inflammatory outcomes during pulmonary Mtb infection, highlighting a role for the endosomal sorting machinery in immunity to intracellular pathogens.

## Introduction

Tuberculosis (TB) remains a significant global health challenge. Although TB-associated mortality declined to 1.23 million people in 2024, TB continued to be the leading cause of infectious disease-related deaths [1]. Incidence increased to 8.3 million cases in 2024, the highest number recorded since global TB monitoring began in 1995 [1]. Despite the identification of *Mycobacterium tuberculosis* (Mtb) nearly 150 years ago, an effective vaccine that reliably prevents adult pulmonary TB is still lacking [2, 3]. Treatment requires prolonged multidrug regimens, and drug-resistant strains continue to emerge [4–6]. These challenges highlight the need to understand how Mtb interacts with host cells, how early immune responses shape pulmonary inflammation, and how these processes influence disease outcome.

Innate immune recognition of Mtb involves pattern recognition receptors that detect bacterial cell wall lipids and nucleic acids. After uptake by macrophages, Mtb resides within membrane-bound phagosomes where it can persist. Macrophage activation facilitates phagosomal maturation via fusion with host endolysosomal compartments and acidification to pH ≤ 5 [7, 8]. However, phagosome maturation and fusion with lysosomes are often disrupted during infection, limiting antimicrobial activity [9–13]. These alterations influence antigen processing and the timing of downstream adaptive immune responses [14–16]. Because protective immunity depends on appropriately coordinated CD4^+^ T cell responses [17–21], the pathways that govern phagosomal maturation in infected phagocytes are critical determinants of host control.

In addition to their essential role in innate cell-autonomous immunity, Mtb-infected macrophages and dendritic cells (DCs) activate adaptive immunity through antigen presentation. Efficient antigen presentation requires lysosomal processing of internalized bacteria, followed by loading of peptide antigens onto MHC class II molecules and transport of peptide-MHC complexes to the cell surface for recognition by T-cell receptors [22]. In humans and mice, initiation of adaptive immunity to Mtb infection is delayed relative to many other infections [23, 24]. Although Mtb is an airway and lung pathogen, antigen-specific CD4^+^ T cell responses are initiated in lung-draining mediastinal lymph nodes, where DCs present processed Mtb antigens to naïve CD4^+^ T cells, leading to their activation, proliferation and migration to the site of infection [25, 26].

Mtb has evolved multiple strategies to limit its exposure to degradative compartments including interference with membrane trafficking and phagosome maturation [10, 11, 27–32] and these mechanisms can reduce antigen processing and prevent efficient antigen presentation [14, 33]. The balance between these bacterial evasion strategies and host endolysosomal processing pathways is a key determinant of whether infection is contained or progresses toward persistent inflammation [34].

Sorting nexins are a family of phosphoinositide-binding proteins that regulate endosomal sorting, membrane remodeling, and intracellular cargo trafficking [35–37]. Sorting nexin 5 (SNX5) contains a phox homology (PX) domain, which allows binding to different phosphoinositides, and a bin, amphiphysin, rvs (BAR) domain that senses and induces membrane curvature and tubulation [38–41]. SNX5 functions as a component of the retromer complex, where it captures endosomal cargo for retrograde trafficking to the Golgi network, thereby influencing the localization and trafficking of multiple cellular components [42]. In addition to its role in the retromer complex, SNX5 associates with SNX4 and SNX17 to form the recycler complex, which mediates the retrieval of autophagy-related proteins, including autophagy-related gene 9 and syntaxin 17, from the autophagosome surface following autophagosome-lysosome fusion [43]. Through these activities, SNX5 is positioned to regulate endolysosomal trafficking events that intersect with both autophagy and antigen processing pathways.

SNX5 has also been implicated in immune cell function. It is required for efficient macropinocytosis by macrophages [44], including uptake of *Mycobacterium bovis* by IFN-γ-activated macrophages [45], and has been linked to antigen processing [44]. These observations suggest that SNX5 may influence how phagocytes acquire and process microbial antigens, although its role in host defense against intracellular bacterial pathogens has not been defined.

Recently, an image-based genome-wide siRNA screen identified SNX5 as a required factor for virus-induced autophagy but not for other forms of stress-induced autophagy [46]. Deletion of *Snx5* increases cellular and murine susceptibility to numerous viruses including Sindbis virus, herpes simplex virus type 1, West Nile virus, and chikungunya virus, resulting in enhanced viral replication [46]. Mechanistically, SNX5 is recruited to virus-containing endosomes, where it promotes the generation of phosphatidylinositol-3-phosphate by the class III phosphatidylinositol-3-kinase complex, thereby activating autophagy-dependent antiviral immune responses [46]. These findings establish SNX5 as a regulator of endosomal signaling events that couple membrane trafficking to immune defense. Although SNX5 has no previously established role in Mtb infection, Mtb similarly traffics through the endolysosomal system following internalization. This parallel raised the possibility that SNX5-dependent trafficking pathways might also influence host responses to intracellular bacterial infection.

Here, we investigated the role of SNX5 during Mtb infection in vivo and in macrophages. We show that in vivo *Snx5* deficiency increased susceptibility to Mtb infection without altering bacterial burden, instead leading to exacerbated lung inflammation. At the cellular level, deletion of *Snx5* did not impair macrophage cell-autonomous antimicrobial immunity but markedly reduced MHC class II-dependent antigen presentation to CD4^+^ T cells. Together, these findings identify SNX5 as a regulator of antigen presentation that shapes inflammatory outcomes during Mtb infection and highlight a previously unappreciated role for an endosomal sorting component in host immunity to tuberculosis.

## Results

### *Snx5*^−/−^ mice succumb earlier than wild-type mice to Mtb infection despite similar organ bacterial burden

To first test if *Snx5* is required for host immune defense during TB, we used an established in vivo model of Mtb infection in which littermate *Snx5*^−/−^ and congenic C57BL/6 mice were infected with a low dose (∼ 200 CFU) of virulent Mtb via aerosol (Figure 1A) [47–51]. Compared to WT mice, *Snx5*^−/−^ mice were more susceptible to Mtb infection, succumbing earlier to disease (Figure 1B). This phenotype was not sex-dependent since both male and female *Snx5*^−/−^ mice showed increased susceptibility to infection (Figure 1B). Surprisingly, lung, liver, and spleen CFU were similar when comparing WT and *Snx5*^−/−^ mice during both early (21 and 42 days post-infection) and chronic stages (90, 120, and 150 days post-infection) of infection (Figure 1C). These results suggest that the absence of SNX5 leads to increased mouse mortality during Mtb infection independent of increased bacterial burden.

**Figure 1:**
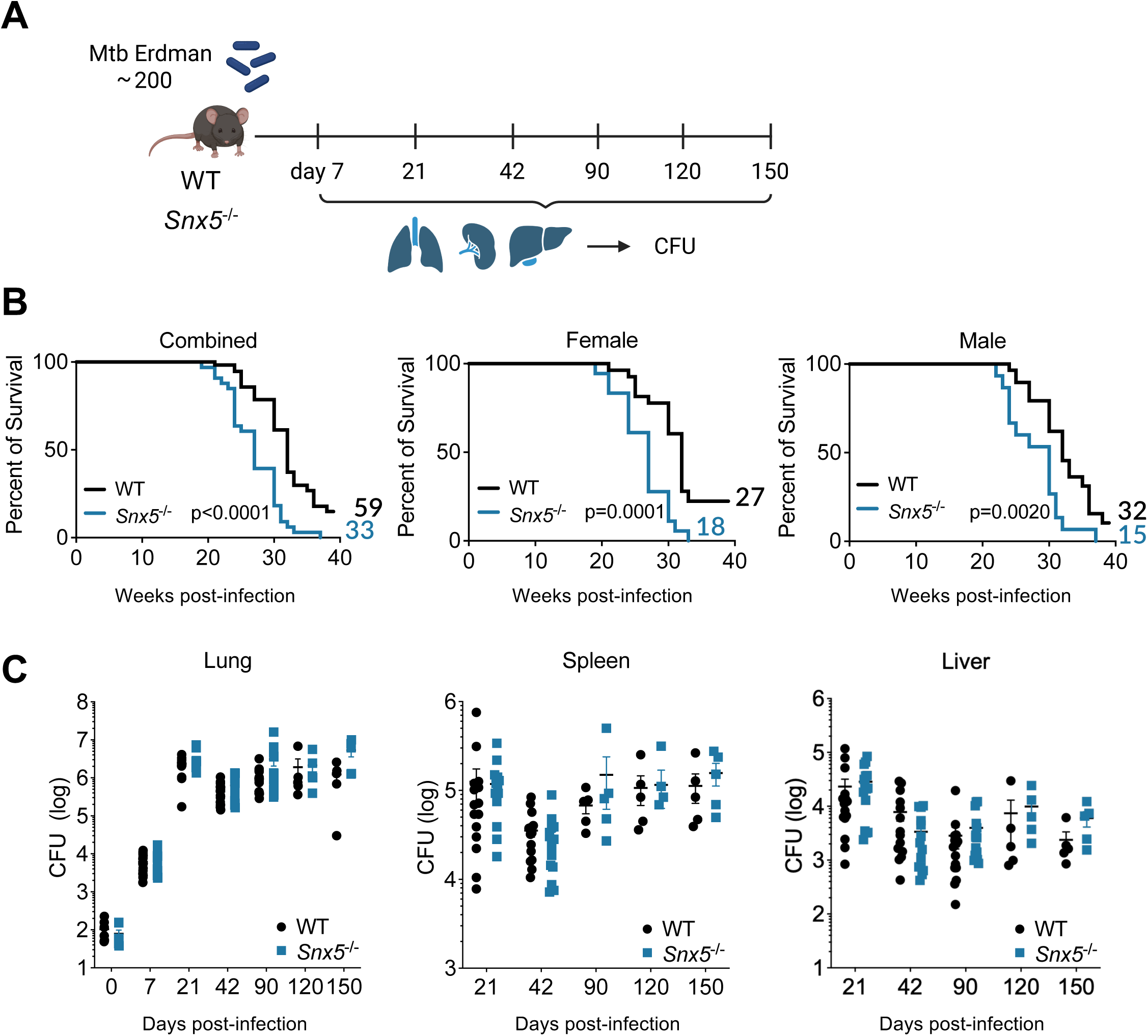
*Snx5*^−/−^ mice succumb to disease earlier than wild-type mice without affecting bacterial replication. (A) WT and *Snx5*^−/−^ mice were infected with approximately 200 CFU of Mtb Erdman. (B) Kaplan-Meier curve of survival of Mtb-infected mice. (C) WT and *Snx5*^−/−^ mice were infected with approximately 200 CFU of Mtb Erdman, and lung, spleen, and liver CFU were determined at 7, 21, 42, 90, 120, and 150 days. (D) Bacterial CFUs in lungs, liver, and spleens of wild-type (WT) or *Snx5*^−/−^ mice at the indicated time point after infection with Mtb. Each symbol corresponds to one animal (Mann-Whitney test).

### Mtb-induced pulmonary inflammation is increased by *Snx5* deletion without increasing the influx of immune cells

Because mouse mortality can be impacted by gene deletion in the absence of differences in Mtb burden [54], a phenotype frequently related to inflammatory immunopathology [55], we analyzed inflammation in lung sections from Mtb-infected wild-type and *Snx5*^−/−^ mice. Using quantification of H&E-stained lung sections from WT and *Snx5*^−/−^ mice (Figure 2A, Supplemental Figure 1), we observed that the percentage of inflamed lungs reached a plateau after 90 days post-infection in WT mice, while inflammatory lung regions in Mtb-infected *Snx5*^−/−^ mice increased over time. At 150 days, lung inflammation was significantly increased in the lungs of *Snx5^-^*^/-^ mice compared to wild-type littermate controls (Figure 2B).

**Figure 2:**
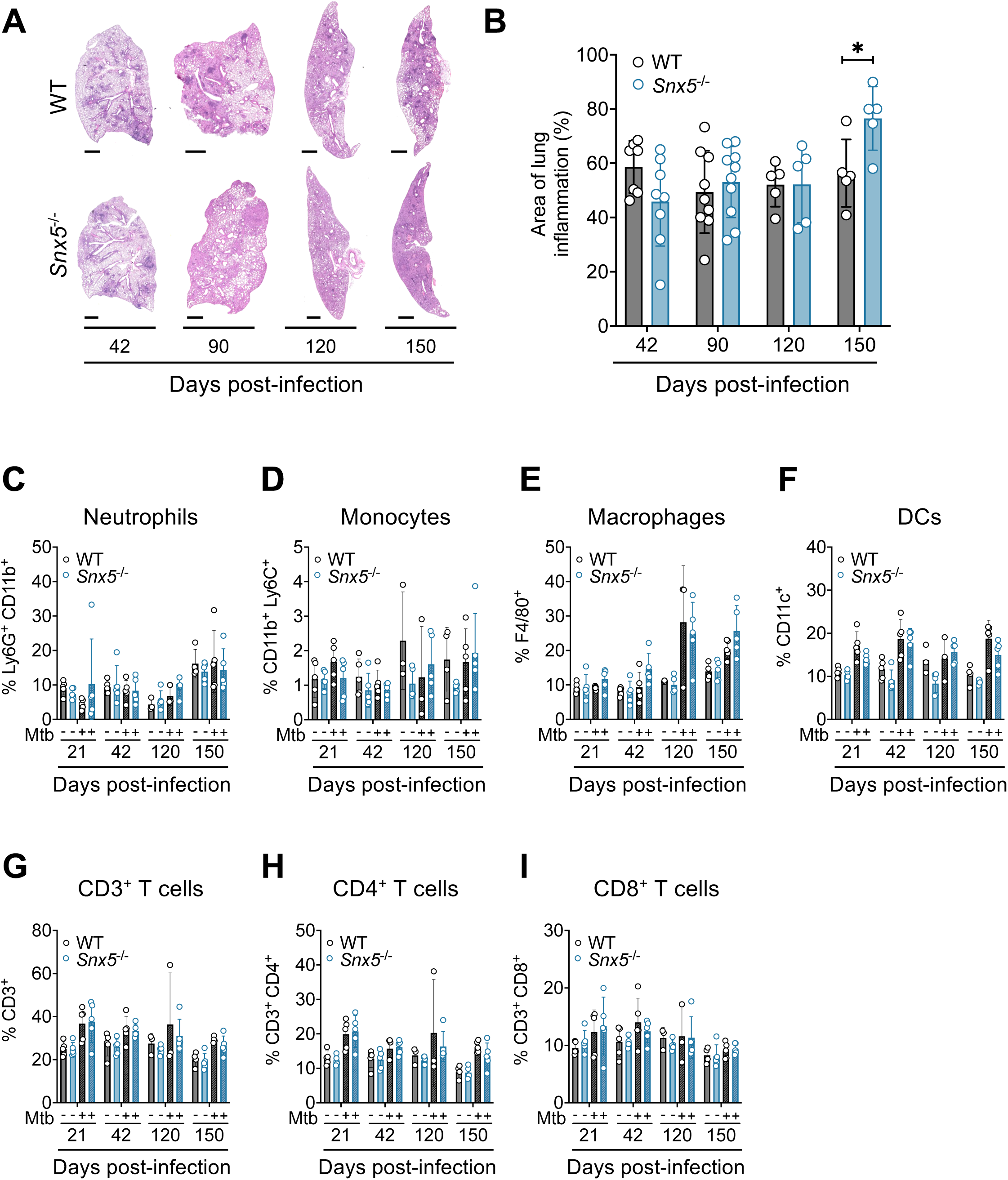
Mtb-induced pulmonary inflammation is increased by *Snx5* deletion without increasing the influx of immune cells. (A) Representative photomicrographs of H&E-stained lungs at 42, 90,120, and 150 days post-infection with Mtb. Scale bars, 1000 μm. (B) Quantitation of inflammatory areas in lungs of Mtb-infected mice at the indicated time point after infection with Mtb. Measurement was performed with ImageJ. Results are the mean ± S.D. for five to ten animals per group. *p < 0.05 (for indicated comparison; Mann-Whitney test). (C-I) Percentage of (C) neutrophils (Ly6G^+^ CD11b^+^), (D) monocytes (CD11b^+^ Ly6C^+^), (E) macrophages (F4/80^+^), (F) DCs (CD11c^+^), (G) T cells (CD3^+^), (H) CD4^+^ T cells (CD3^+^ CD4^+^) and (I) CD8^+^ T cells (CD3^+^ CD8^+^) in the lung of mice during low-dose Mtb infection. Each circle corresponds to one animal. Bars are the mean ± S.D. for 3 to 5 animals per group per time point. Statistical analysis was by one-way ANOVA.

To investigate whether the increased inflammation in the lung of *Snx5*^−/−^ mice was due to increased lung-infiltrating populations during Mtb infection, lung cells from WT and *Snx5*^−/−^ mice infected or not with Mtb were analyzed by flow cytometry using myeloid and lymphocyte markers. There was no difference in the influx of neutrophils (Ly6G^+^ CD11b^+^), monocytes (CD11b^+^ Ly6C^+^), macrophages (F4/80^+^), DCs (CD11c^+^), or T cells (CD3^+^) into the lungs of WT and S*nx5*^−/−^ mice (Figure 2C-I, Supplemental Figure 2).

Because we did not observe a difference in the influx of CD3^+^ T cells to the lungs, we next investigated whether *Snx5* deletion impacted the ability of T cells to produce pro-inflammatory cytokines. We first measured cytokine levels in lung homogenates of mice. No difference was observed in the production of IFN-γ, TNF-α, or IL-2 (Figure 3A-C). However, *Snx5* deficiency was associated with decreased production of IL-12p40 in the lungs of Mtb-infected mice (Figure 3D). Next, to determine if production of cytokines differed in explanted T-cells, ex vivo lung cells, and splenocytes activated with CD3 and CD28 were processed for intracellular staining and analyzed by flow cytometry to characterize cytokine production by CD4^+^ T cells (Supplemental Figure 3). As expected, the production of IFN-γ and TNF-α was increased in lung CD4^+^ T cells from mice infected with Mtb (Figure 3E and 3F). However, production of IFN-γ, TNF-α, and IL-2 was comparable between lung CD4^+^ T cells from WT and *Snx5*^−/−^ mice (Figure 3E-G). Similar results were observed in ex vivo splenocyte derived CD4^+^ T cells from WT and *Snx5*^−/−^ mice (Extended Data Fig. 4). Together, these findings indicate that *Snx5* deletion is associated with increased lung inflammation during Mtb infection without altering immune cell recruitment or CD4^+^ T-cell cytokine production, suggesting a role for SNX5 in limiting inflammation-driven immunopathology.

**Figure 3:**
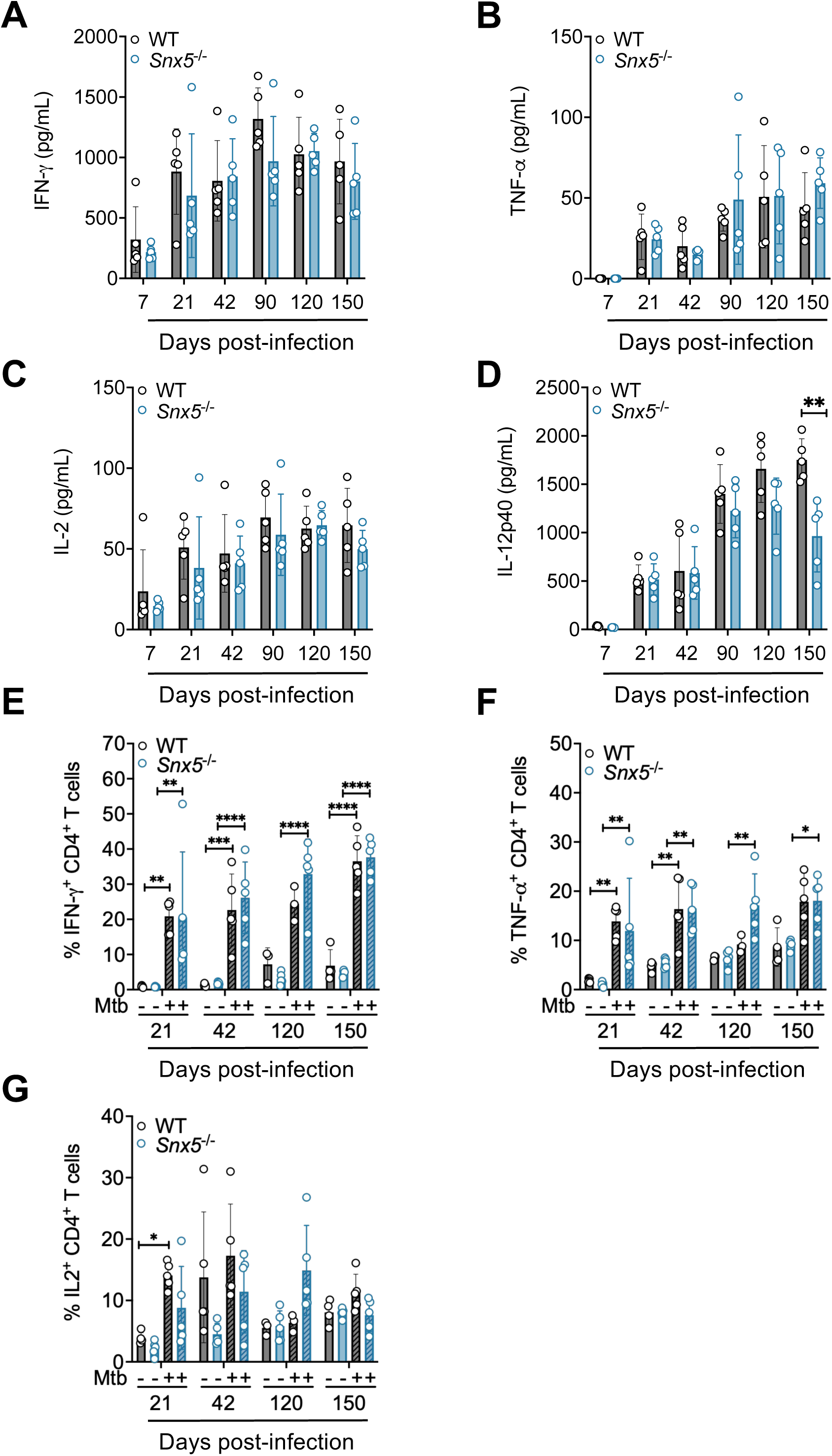
*Snx5* deficiency decreases the production of IL-12p40 in lung homogenates of Mtb-infected mice. (A) IFN-γ, (B) TNF-α, (C) IL-2, and (D) IL-12p40 were assessed in supernatants of homogenized lungs by ELISA. **p < 0.01 (for indicated comparison; Mann-Whitney test). (E-G) Lung explanted CD4^+^ T cells were characterized for cytokine expression, including (E) IFN-γ, (F) TNF-α, and (G) IL-2 using surface and intracellular cytokine staining for flow cytometry. Each circle corresponds to one animal. Bars are the mean ± S.D. for 3 to 5 animals per group per time point. *p < 0.05, **p < 0.01, ***p < 0.001 and ****p < 0.0001 (for indicated comparison; one-way ANOVA)

### *Snx5* deletion does not impact macrophage cell-autonomous immunity during Mtb infection

Since the role of SNX5 during Mtb infection has not been previously explored, we next assessed whether Mtb modulates SNX5 expression in BMDMs. At 4 and 24 h post-infection, no significant changes in SNX5 expression were observed in either resting or IFN-γ-activated macrophages (Supplemental Figure 5A and B). We also investigated whether *Snx5* deletion alters the expression of SNX6 and SNX32, closely related members of the SNX-BAR family known for their high homology with SNX5 and interchangeable roles [56]. We observed that in BMDM from *Snx5*^−/−^ mice (Figure 4A), *Snx5* deletion did not impact the expression of SNX6 or SNX32 at 24 h post-infection (Supplemental Figure 5C-E).

**Figure 4:**
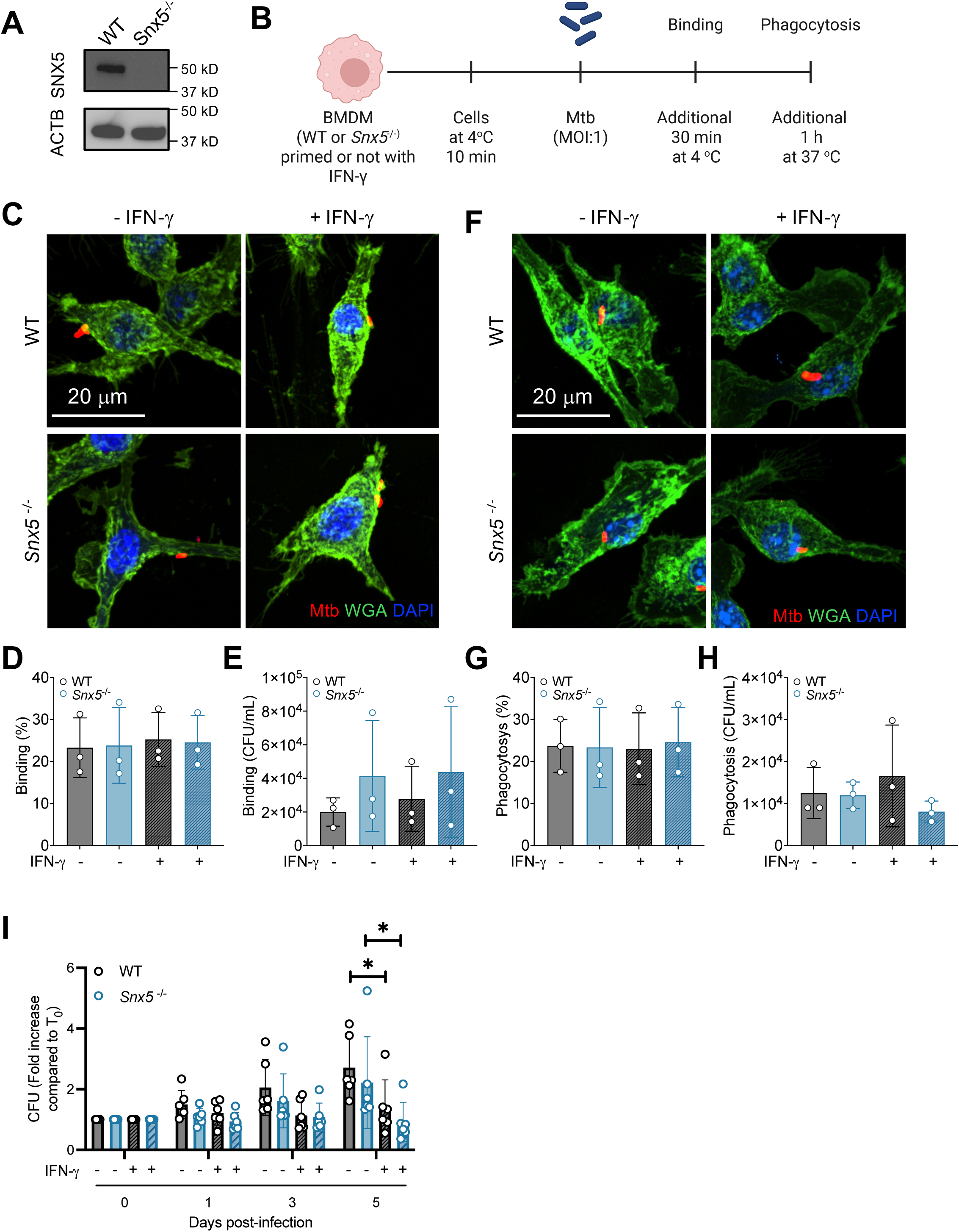
*Snx5* deletion does not impact macrophage cell-autonomous role in controlling Mtb infection. (A) Representative western blot to detect SNX5 levels in BMDMs from wild-type (WT) and *Snx5*^−/−^ mice. ACTB: Actin. (B) Description of the binding and phagocytosis experimental procedures. (C-H) Wild-type and *Snx5* ^−/−^BMDMs either primed or not with IFN-γ, were infected with mCherry-Mtb (red). (C-E) Binding and (F-H) phagocytosis were analyzed by confocal microscopy with quantification of bacterial number and by quantification of bacterial CFU. For confocal microscopy, cell nuclei were labeled with DAPI (blue). Scale bars, 20 µm. Bars are the mean ± S.E.M. from three independent experiments. The circles shown correspond to each experiment. (I) Mycobacterial growth in wild-type or *Snx5*^−/−^ BMDMs infected with Mtb, either primed or not with IFN-γ. Bars are the mean ± S.E.M. from three independent experiments. The circles shown correspond to each experiment *p < 0.05 (for indicated comparison; two-way ANOVA).

To test whether *Snx5* has a cell-autonomous role in controlling Mtb infection, we infected BMDMs from wild-type or *Snx5*^−/−^ mice and quantified mycobacterial binding, phagocytosis, and intracellular growth. To investigate the impact of *Snx5* deletion on Mtb binding, BMDMs were kept at 4 °C, allowing bacteria-cell interaction without bacterial phagocytosis (Figure 4B). No difference in Mtb binding was observed between BMDMs from wild-type or *Snx5*^−/−^ mice, either primed or not primed with IFN-γ (Figure 4C-E). Similarly, *Snx5* deletion did not impact Mtb phagocytosis by BMDMs (Figure 4F-H). Additionally, after 5 days of infection, IFN-γ-activated BMDMs from wild-type or *Snx5*^−/−^ mice controlled Mtb replication to a similar extent (Figure 4I).

Due to the role of SNX5 in membrane trafficking [57], we next evaluated the impact of SNX5 on the maturation of the Mtb-containing vacuole. We analyzed colocalization of Mtb with markers of early endosomes (EEA1) and lysosomes (LAMP1) using immunofluorescence microscopy of fixed cells. Specifically, we infected WT and *Snx5*^−/−^ macrophages with mCherry-Mtb and fixed cells 24 h after infection. As shown in Figure 5, the recruitment of EEA1 (Figure 5A and 5B) and LAMP1 (Figure 5C and 5D) to Mtb-containing vacuoles was similar in *Snx5*^−/−^ and wild-type macrophages, despite priming with IFN-γ. These results suggest that deficiency of *Snx5* does not impact macrophage cell-autonomous immunity during Mtb infection.

**Figure 5:**
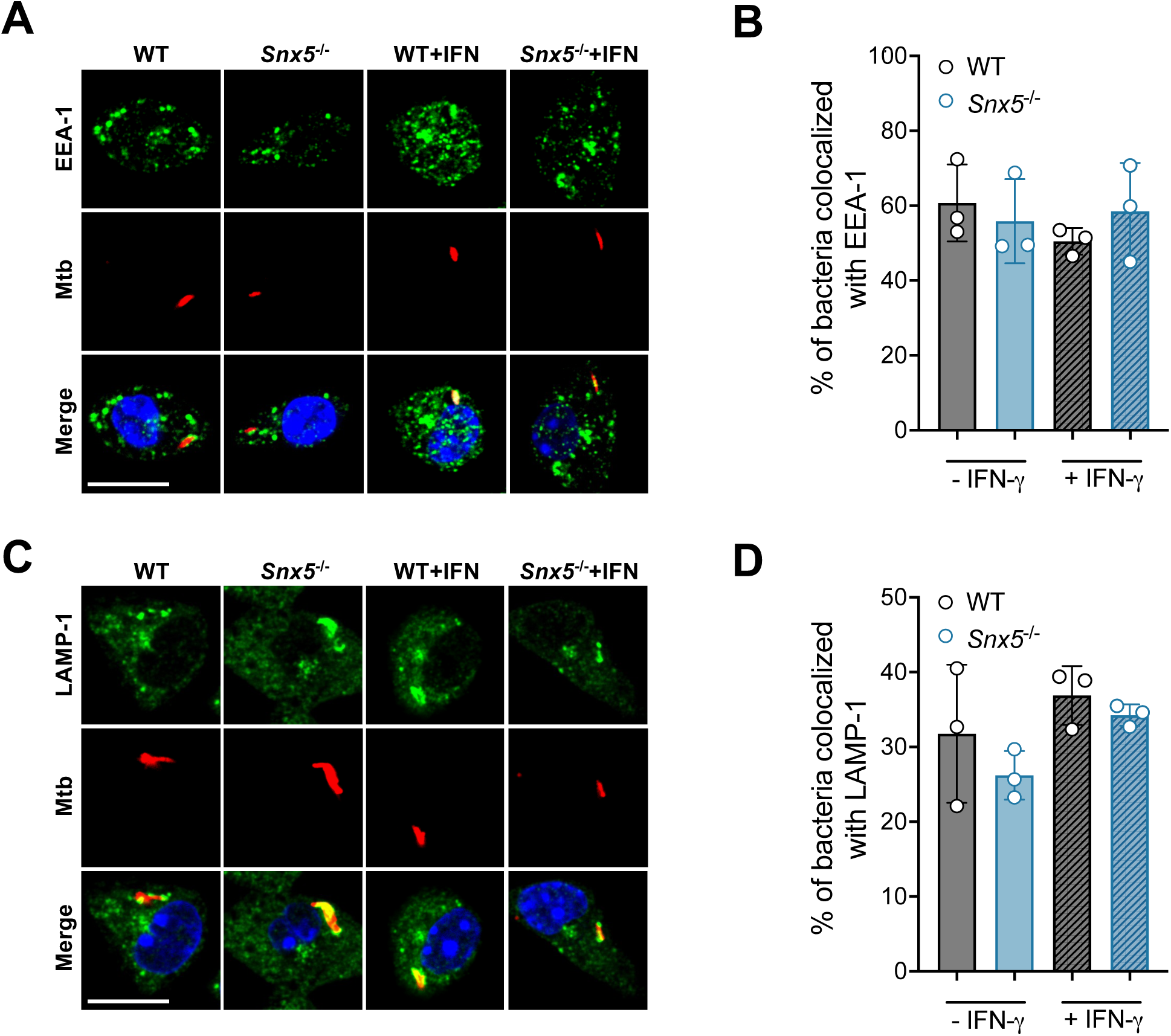
*Snx5* deletion does not alter the recruitment of EEA1 and LAMP1 to the Mtb-containing vacuole. WT and *Snx5*^−/−^ BMDMs, either primed or not with IFN-γ, were infected with mCherry-Mtb (red). After 24 h, cells were fixed and stained for (A and B) EEA1 or (C and D) LAMP1 (green). For confocal microscopy, cell nuclei were labeled with DAPI (blue). Representative photomicrographs (A and C) and quantitation (B and D) of the colocalization of mCherry-Mtb and EEA1 (A and B), or LAMP1 (C and D). For each coverslip, at least 100 bacteria were evaluated. Scale bars, 10 μm. Bar graphs are the mean ± S.D. from one experiment performed in quadruplicate. The circles shown correspond to each replica. Statistical analysis was by t-test.

### SNX5 does not affect macrophage transcriptional responses to Mtb infection

To understand how SNX5 might impact mouse susceptibility to Mtb infection, we used bulk RNA sequencing to investigate its impact on macrophage transcriptional responses to Mtb infection. To that end, we compared the transcriptomes of WT and *Snx5*^−/−^ BMDMs, uninfected or Mtb-infected, at 4 and 24 h post-infection. Principal component analysis (PCA) clearly separated infected and uninfected cells. However, it does not distinguish between WT and *Snx5*^−/−^ cells (Supplemental Figure 6A). Genome mapping identified 152 and 187 significantly upregulated genes and 4 and 41 significantly downregulated genes in the WT-infected cells compared to uninfected WT cells after 4 and 24 h of infection, respectively (Supplemental Figure 6B). In Mtb-infected *Snx5*^−/−^ cells, 208 and 300 genes were upregulated, while only 2 and 55 were downregulated after 4 and 24 h of infection, respectively, compared to uninfected *Snx5*^−/−^cells (Supplemental Figure 6B).

As expected, in both WT and *Snx5*^−/−^ murine BMDMs infected with Mtb, we observed upregulation of classical immunologically significant genes. These included several chemokines (e.g., CCL2, CCL3, CCL4, CCL5), cytokines (e.g., IL1A, IL1B, IL6, TNF), signaling molecules (e.g., TNIP3, NOD2, SQSTM1 (p62), NLRP3), interferon-stimulated genes (e.g., IFIT1, ISG20, IGTP, IIGP1), and transcription factors (e.g., IRF1, IRF7, STAT1, STAT2) (Supplemental Figure 6C). Gene Ontology (GO) biological process enrichment analysis further revealed that among upregulated genes were genes with described roles in responses to cytokines, defense responses, and responses to external biotic stimuli (Supplemental Figure 6D). However, consistent with the PCA analysis, macrophage transcriptomes of *Snx5*^−/−^ and wild-type BMDMs, either infected or not with Mtb, did not demonstrate any meaningful differences between the two genotypes (Supplemental Figure 6D). Only one differentially expressed gene (DEG) was identified between *Snx5*^−/−^ and wild-type BMDMs at 4 h (CRYAB, downregulated in *Snx5*^−/−^ uninfected cells) and 24 h (CAPN1, upregulated in *Snx5*^−/−^ infected cells) after infection (Supplemental Figure 6B). Thus, we conclude that *Snx5* does not affect the transcriptional response of macrophages to Mtb infection.

### *Snx5* modestly affects pro-inflammatory cytokine production by BMDMs

Since we observed that *Snx5* deletion decreased IL12p40 production in lung homogenates of Mtb-infected mice, we next assessed if the production of pro-inflammatory cytokines by Mtb-infected macrophages in vitro is affected by *Snx5* deletion (Figure 6). In IFN-γ-activated, but not resting, macrophages, *Snx5* deletion resulted in increased TNF-α secretion (Figure 6A) and, consistent with our in vivo study, decreased IL-12p40 secretion (Figure 6B). No differences were observed in the production of MCP-1 or IL-6 (Figure 6C and 6D).

**Figure 6:**
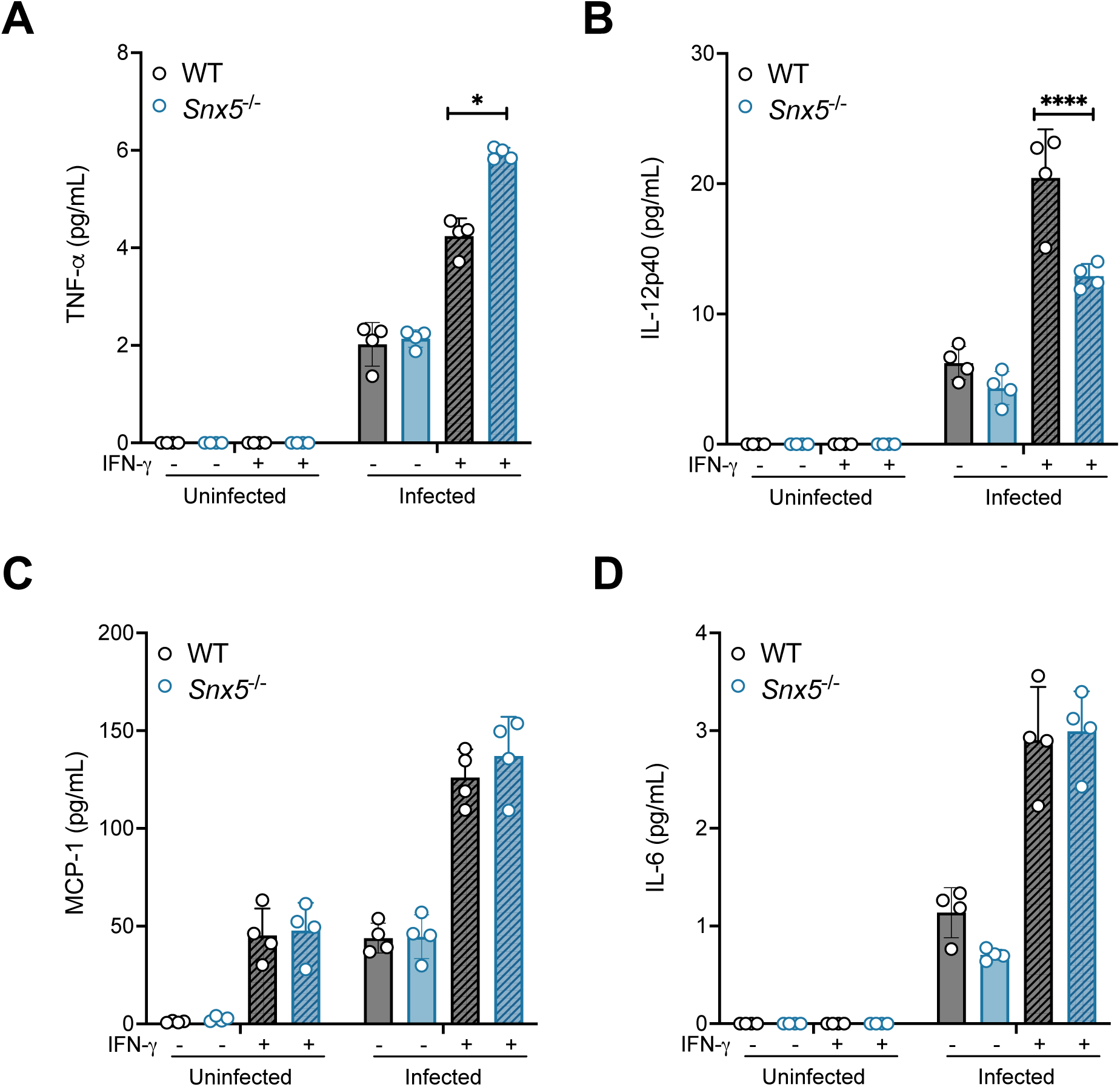
Lack of SNX5 modestly affects pro-inflammatory cytokine production by BMDM. (A) TNF-α, (B) IL-12p40, (C) MCP-1, and (D) IL-6 were assessed in cell supernatants of infected macrophages, either primed or not with IFN-γ (100 UI/mL). Bars are the mean ± S.D. from one experiment performed in quadruplicate. The circles shown correspond to each replica. *p < 0.05 and ****p < 0.0001 (for indicated comparison; one-way ANOVA).

### *Snx5* is required for efficient antigen processing and MHC class II-restricted antigen presentation by macrophages and DCs

Since *Snx5* has previously been shown to impact antigen processing [44] and antigen presentation in B cells [58], we hypothesized that *Snx5* is required for antigen presentation during Mtb infection. We first assessed whether *Snx5* deficiency impacts antigen processing in our model. To this end, resting or IFN-γ-activated WT and *Snx5* ^−/−^BMDMs were infected or not with Mtb and incubated with DQ-BSA, a marker of degradative compartments. Consistent with a prior report [44], *Snx5* deletion reduced DQ-BSA intensity in both Mtb-infected and uninfected BMDMs, regardless of IFN-γ priming (Figure 7A and 7B). However, no difference was observed in the colocalization of Mtb and DQ-BSA between WT and *Snx5* ^−/−^ infected cells (Figure 7C).

**Figure 7:**
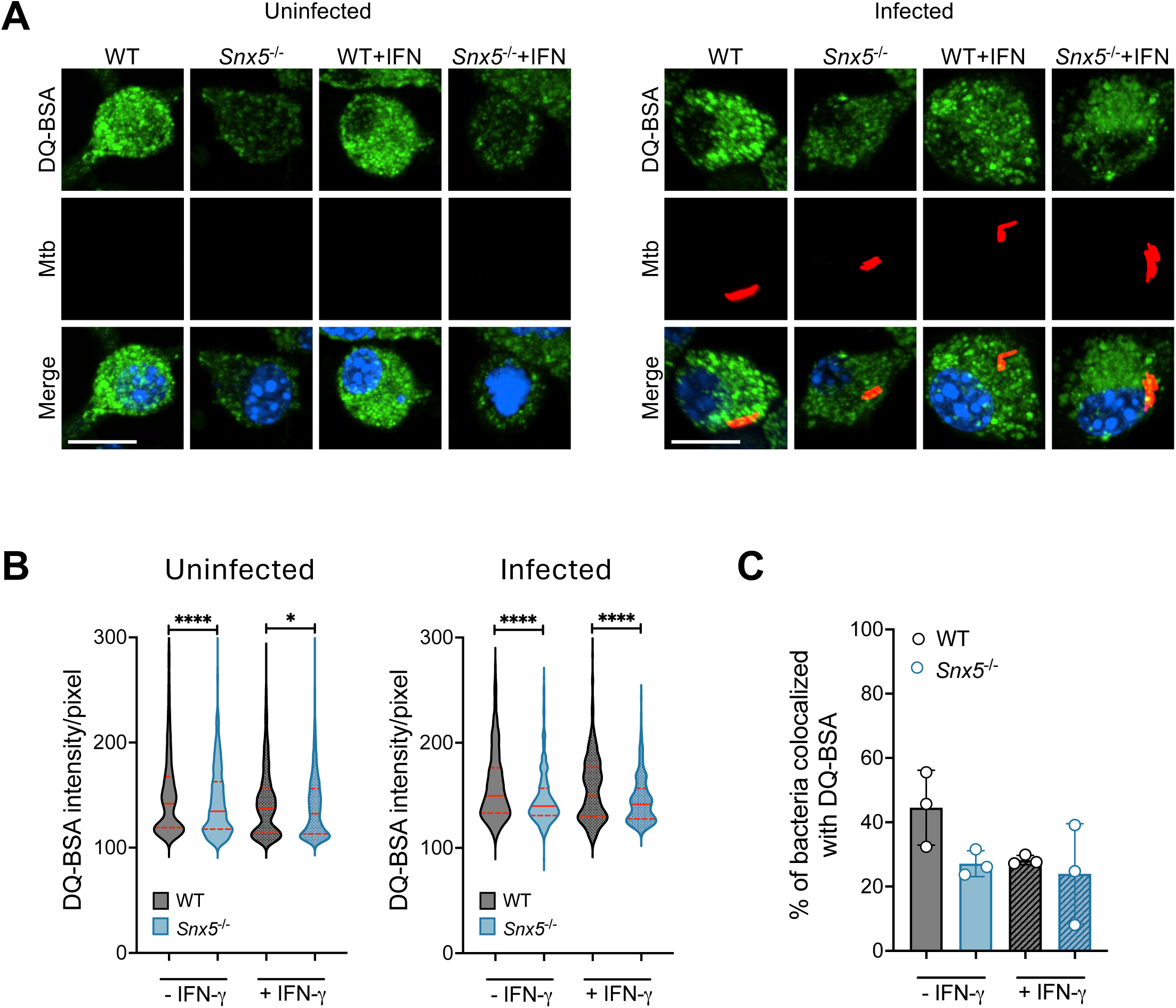
*Snx5* deletion mildly reduced DQ-BSA intensity in macrophages. (A-C) WT and *Snx5*^−/−^ BMDMs, either primed or not with IFN-γ, were infected or not with mCherry-Mtb (red). After 24h, cells were incubated with DQ-BSA (green) for 30 min and then fixed. For confocal microscopy, cell nuclei were labeled with DAPI (blue). (A) Representative photomicrographs. Scale bars, 10 μm. (B) Quantitation of DQ-BSA intensity per pixel. For each coverslip, at least 50 cells were evaluated. Lines represent the median and violin plot quartiles (25% and 75%) *p < 0.05, ****p < 0.0001 (for indicated comparison; Mann-Whitney test). (C) Quantification of the colocalization of mCherry-Mtb and DQ-BSA. For each coverslip, at least 100 bacteria were evaluated. Bars are the mean ± S.D. from one of three experiments performed in triplicate. The circles shown correspond to each replica. Statistical analysis was by Mann-Whitney test.

We next evaluated the functional consequences of *Snx5* deletion on antigen presentation. We incubated macrophages with the known T cell epitope OVA_323-339_ [59, 60], the full-length Mtb protein Ag85B [15], or heat-killed Mtb and then co-cultured them with CD4^+^ T cell hybridoma cells with specificity against either OVA_323-339_ (DO11.10) [59, 60] or Ag85B (BB7) [15]. We then quantified IL-2 accumulation in the supernatant as a measure of CD4^+^ T cell activation [59, 61] (Figure 8A). Deletion of *Snx5* in BMDMs resulted in reduced IL-2 production by MHC class II-restricted CD4^+^ T cells when IFN-γ-stimulated macrophages were incubated with full-length Ag85B (Figure 8B, Supplemental Figure 7) and heat-killed Mtb (Figure 8C, Supplemental Figure 7).

**Figure 8:**
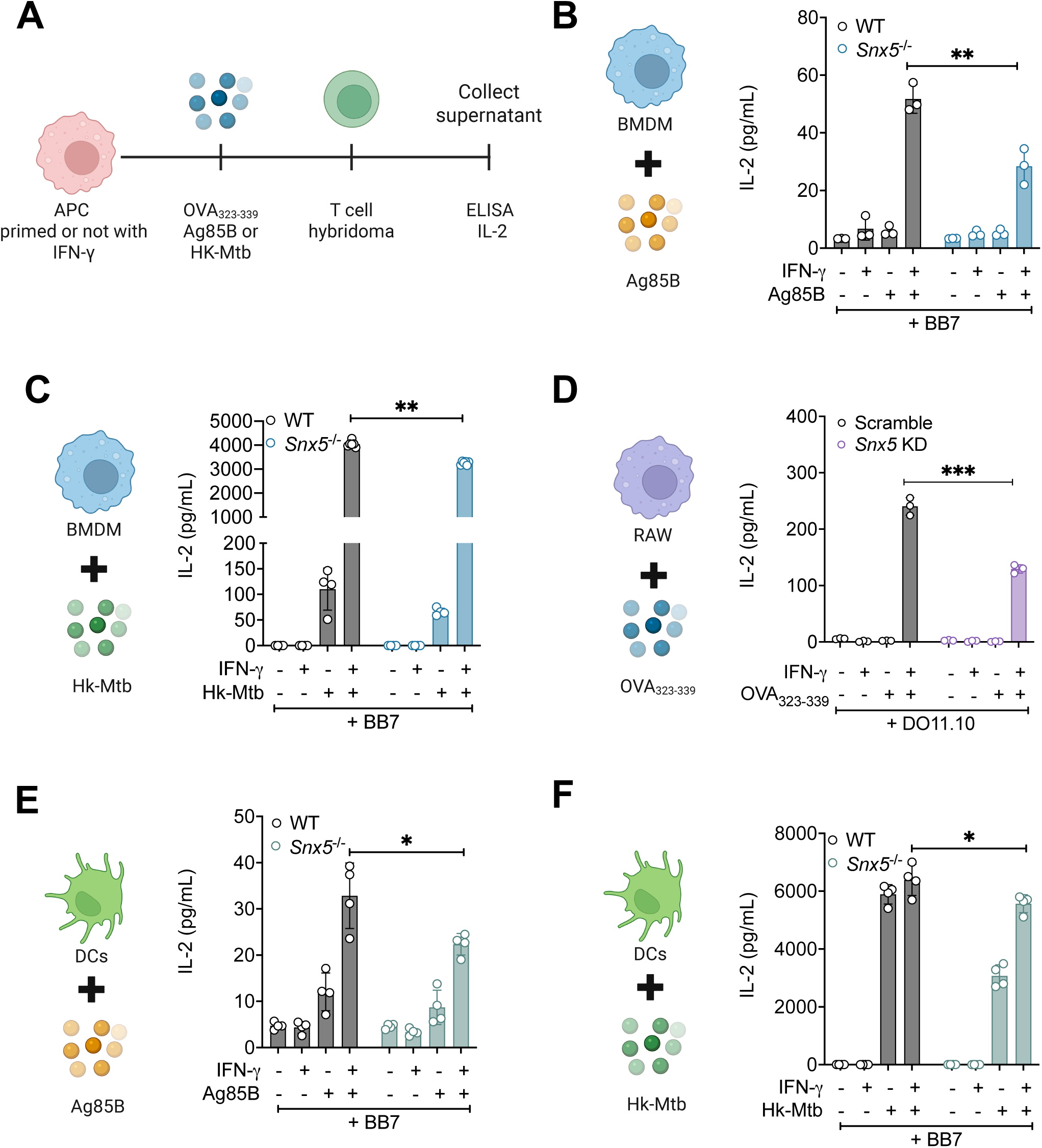
SNX5 is necessary for efficient antigen presentation by macrophages and DCs. (A) Timeline of antigen presentation assay. (B) RAW 264.7, (C and D) BMDMs or (E and F) DCs, either primed or not with IFN-γ, were fed with OVA _323-339_, Mtb Ag85B, or heat-killed Mtb as depicted on the graphs. After (B) 4 or (C-F) 18h, the cells were incubated with antigen-specific MHC class II-restricted T cell hybridomas. After 24h, supernatants were assessed for IL-2 secretion by T cells. Bars are the mean ± S.D. from one of three experiments performed at least in triplicate. The circles shown correspond to each replica. *p < 0.05, **p < 0.01 and ***p < 0.001 (for indicated comparison; t-test).

Importantly, this reduction was not due to an impaired ability of macrophages to process MHC class II antigens, as similar results were observed when RAW 264.7 cells were fed the mature MHC class II peptide OVA_323-339_ (Figure 8D, Supplemental Figure 7). Because dendritic cells are the most efficient antigen-presenting cells, we next investigated whether *Snx5* was also required for antigen presentation by DCs. As expected, DCs did not require IFN-γ to induce efficient antigen presentation, and like macrophages, *Snx5* deletion reduced antigen presentation by approximately 40% (Figure 8E and F, Supplemental Figure 7).

### *Snx5* promotes peptide loading onto MHC class II without altering surface expression of antigen presentation machinery

Expression of peptide-MHC complexes on the surface of antigen-presenting cells is essential for efficient antigen presentation to T cells. To determine whether SNX5 deletion affects the MHC class II antigen presentation machinery, we assessed the expression of MHC class II and the costimulatory molecules, CD80 and CD86, on the surface of *Snx5*^−/−^ and wild-type BMDMs. No differences were observed in the cell surface expression of MHC-II (Figure 9A, Supplemental Figure 8A), CD80 (Figure 9B, Supplemental Figure 8B), and CD86 (Figure 9C, Supplemental Figure 8C) comparing *Snx5*^−/−^ and WT BMDMs.

**Figure 9:**
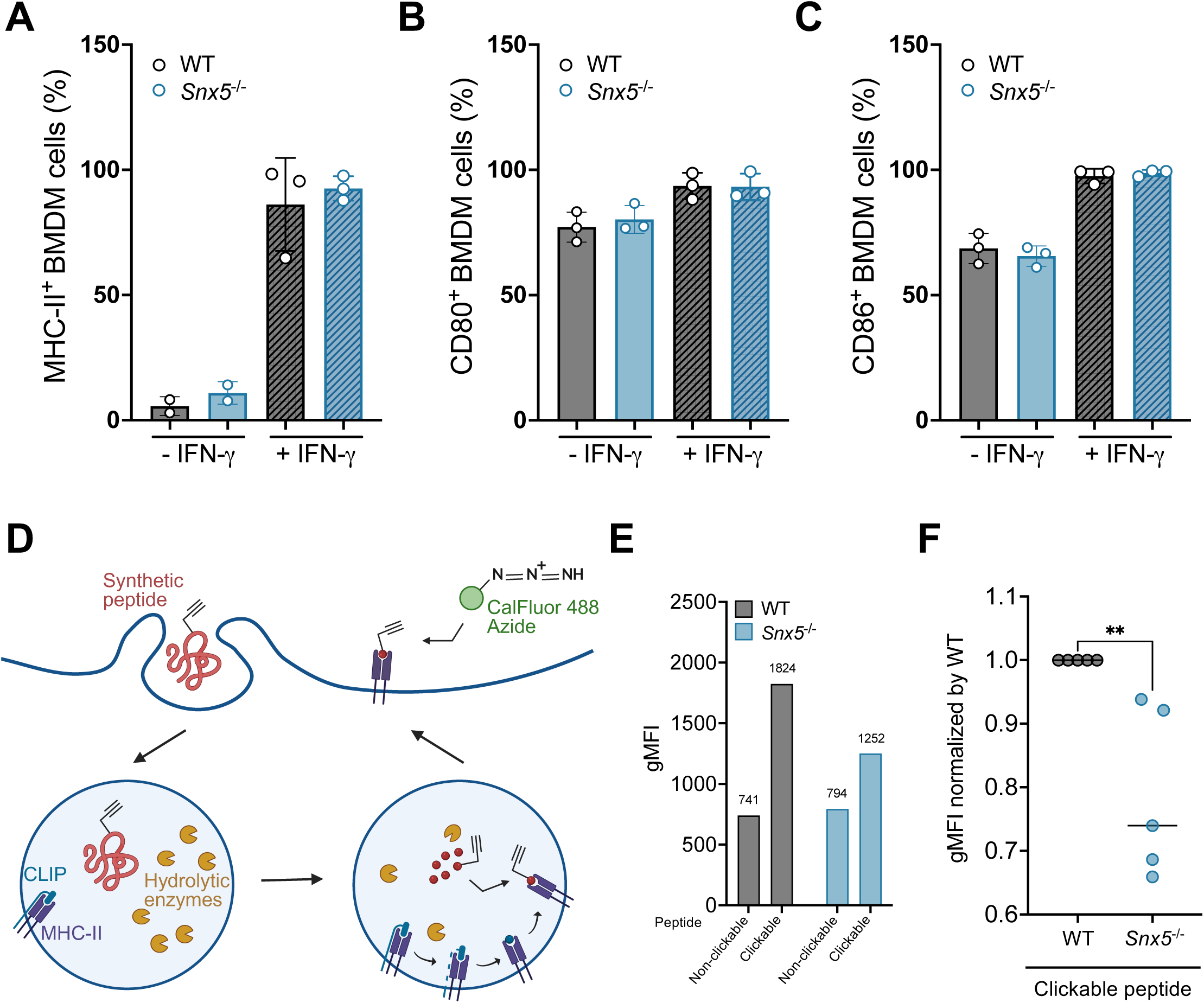
*Snx5* deletion reduces virus HA peptide loading onto MHC class II without affecting surface expression of MHC-II and costimulatory molecules. (A-C) Expression of (A) MHC class II, (B) CD80, and (C) CD86 molecules on the surface of wild-type or *Snx5*^−/−^ BMDMs, either primed or not with IFN-γ (100 UI/mL), analyzed by flow cytometry. Bars are the mean ± S.D. from three independent experiments. The circles shown correspond to each experiment. (D-E) IFN-γ-activated BMDMs were incubated with HA peptides for 18 h and then labeled with CalFluor 488. (D) Representative figure of the click chemistry assay to detect HA-peptide loaded on MHC-II, adapted from [52]. BMDMs were incubated with the extended influenza A virus hemagglutinin (HA) peptide (HA_318-338_) containing a non-naturalized amino acid (a “clickable” peptide) for 18 h. This peptide must be cleaved and loaded onto MHC-II for presentation on the cell surface, where it can be detected by CalFluor 488. (F) Representative quantification of HA-CalFluor-488 quantification (gMFI – geometric mean fluorescence intensities). Bars are the mean ± S.D. from one of five independent experiments. (G) Quantification of HA-CalFluor-488 (gMFI) normalized to control WT cells. Circles represent individual experiments. **p < 0.01 (for indicated comparison; t-test).

We therefore hypothesized that the reduced T cell activation observed in *Snx5*^−/−^BMDMs was associated with impaired peptide loading to MHC-II. To test this, we adapted a protocol whereby cells are incubated with an HA-peptide containing an unnatural amino acid, propargylglycine, that permits detection of surface-exposed peptide-MHC class II complexes via click chemistry to a fluorescent dye (Figure 9D) [52]. In WT control cells incubated with the non-clickable HA_318-338_ peptide (lacking propargylglycine), the CalFluor 488 signal was approximately two-fold lower compared to the cells incubated with the clickable peptide (Figure 9E, Supplemental Figure 8D). Notably, CalFluor background labeling did not differ between WT and *Snx5*^−/−^ BMDMs.

Among cells incubated with the clickable peptide, we observed a reduction of approximately 30% in CalFluor 488 signal in *Snx5*^−/−^ BMDMs compared to WT controls (Figure 9E and F, Supplemental Figure 8D). Taken together, these data indicate that SNX5 is required for efficient peptide loading onto MHC class II molecules, resulting in reduced levels of peptide-loaded MHC-II at the cell surface without altering total surface expression of MHC-II or costimulatory molecules.

## Discussion

In this study, we identify SNX5 as an important regulator of host defense during Mtb infection. Although *Snx5* deficiency did not alter bacterial burden, *Snx5*^−/−^ mice were more susceptible to Mtb infection, a phenotype associated with exacerbated pulmonary inflammation. At the cellular level, we show that SNX5 is required for efficient antigen presentation to MHC class II-restricted CD4^+^ T cells by both macrophages and dendritic cells, without affecting total MHC-II or costimulatory molecule surface expression, but instead reducing the amount of peptide-loaded MHC-II on the surface of antigen-presenting cells. Together, our findings reveal a previously unappreciated role for SNX5 in regulating Mtb immune responses and inflammation-driven immunopathology.

The increased mortality of *Snx5*^−/−^ mice in the absence of higher bacterial loads suggests that factors beyond bacterial replication contribute to disease progression. Similar phenotypes have been observed in other models of Mtb infection, where loss of disease tolerance rather than pathogen burden drives mortality [54, 62, 63]. For example, deletion of *Cybb* (gp91), a subunit of the PHOX multi-protein complex, leads to increased mouse mortality due to excessive lung inflammation caused by increased caspase-1 activation and neutrophil infiltration [63]. Likewise, *Ppif* (Cyclophilin D) knockout mice exhibit increased T cell proliferation associated with enhanced T cell metabolic activity [54]. In our study, histological analysis revealed that *Snx5*^−/−^ mice exhibited more extensive lung inflammation than WT control mice, consistent with an inability to balance protective and pathological immune responses. Notably, the exacerbated inflammation was not associated with increased influx of immune cells into the lung, and we observed only mild effects on pro-inflammatory cytokine production in lung homogenates. Thus, further studies will be required to elucidate how *Snx5* regulates lung inflammation during Mtb infection. Because we used whole-body knockout mice, it is also possible that non-immune cells contributed to the phenotype reported here.

At the cellular level, we found that *Snx5* has no impact on macrophage cell-autonomous response against Mtb. This contrasts with previous work showing that depletion of SNX5/6 increases *Chlamydia trachomatis* progeny production in HeLa cells [64]. In the context of viral infection, SNX5 restricts infection by several viruses (Sindbis virus, herpes simplex virus type 1, West Nile virus, and chikungunya virus) in both HeLa cells and primary mouse embryonic fibroblasts via an autophagy-dependent mechanism [46]. Together, these findings suggest that SNX5’s role during infection may be both pathogen– and cell type-specific, likely reflecting differential engagement of membrane trafficking and autophagic pathways by distinct intracellular pathogens.

Our finding that SNX5 promotes antigen presentation adds to growing evidence implicating SNX family members in endomembrane trafficking and immune signaling [65–69]. Previous studies showed that SNX5 supports antigen processing in macrophages [44] and participates in antigen presentation in B cells by coordinating actin remodeling and lysosomal dynamics [58]. Consistent with these reports, we observed that *Snx5* deficiency mildly reduced degradation of DQ-BSA, suggesting impaired endolysosomal proteolysis, and decreased peptide-loaded MHC-II at the cell surface. Importantly, the lack of changes in the total MHC-II and costimulatory molecule expression indicates that SNX5 specifically affects the generation and/or trafficking of peptide-MHC-II complexes rather than their transcriptional regulation. This interpretation is further supported by our transcriptomic analysis, which revealed no differences in gene expression between *Snx5*-knockout and WT BMDM controls.

Successful antigen presentation depends on multiple coordinated events, ranging from antigen acquisition to the display of peptide-MHC complexes on the cell surface [70]. Based on our data, SNX5 may facilitate antigen presentation through multiple, non-mutually exclusive pathways. First, SNX5 may impact peptide loading within the MHC-II compartment, potentially by regulating endosomal-lysosomal dynamics required for optimal processing and loading, consistent with our observed proteolytic activity defects and prior work [44]. Second, SNX5 may affect post-loading trafficking of peptide-MHC-II complexes to the plasma membrane, as SNX5 has been shown to participate in the recycling and retrograde trafficking of several membrane receptors [38, 39, 71, 72]. Whether SNX5’s effect on antigen presentation depends on its association with the retromer complex or reflects a retromer-independent role remains an open question.

As previously noted, we observed increased susceptibility of *Snx5^−/−^*mice to Mtb infection, associated with higher pulmonary inflammation that was not accompanied by increased influx of neutrophils, macrophages, DCs, monocytes, or T cells. A recent study demonstrated that the spatial organization of inflammatory lesions, rather than cell numbers, can dramatically influence disease outcomes during Mtb infection [73]. In this context, combining our in vivo findings with the observed reduction in antigen presentation in *Snx5^−/−^* cells raises the possibility that SNX5 may influence the spatial organization or local interactions of immune cells in the lung, thereby shaping the quality and localization of inflammatory response against Mtb.

A limitation of our study is that we did not establish the molecular mechanism by which SNX5 regulates peptide loading and presentation. Although the clickable-peptide assay indicates reduced surface peptide-MHC-II complexes, it remains unclear whether this defect arises from peptide exchange dynamics, changes in vesicular trafficking, or both. In addition, because this assay used an influenza A-derived peptide, developing Mtb-specific clickable peptides will be necessary to determine whether SNX5 selectively affects the presentation of particular mycobacterial epitopes or more broadly impacts antigen presentation. Furthermore, we did not directly assess whether SNX5 impacts T-cell priming in vivo or how this relates to the increased mortality in *Snx5*-deficient mice, which remain important questions for future investigation.

In conclusion, we demonstrate that SNX5 is a key regulator of the host immune response to Mtb, promoting efficient antigen presentation and controlling inflammation without affecting bacterial burden. These findings extend the known functions of *Snx5* beyond endosomal trafficking and antiviral defense to include regulation of antibacterial immunity and immunopathology.

## Material and Methods

### Mice

*Snx5*^−/−^ mice, backcrossed to the C57BL/6J genetic background for more than ten generations, were obtained from Beth Levine [46] and bred to C57BL/6J mice (The Jackson Laboratory) to obtain *Snx5*^+/−^ heterozygotes. Littermate wild-type controls produced by heterozygous crosses were used for all experiments. All experimental protocols involving the use of mice were approved by the Institutional Animal Care and Use Committee (IACUC) of the University of Texas (UT) Southwestern Medical Center (protocol number 2017-101977).

### *Mycobacterium tuberculosis* (Mtb) strains

Mtb Erdman wild-type and mCherry-expressing Mtb (mCherry-Mtb) strains were cultivated in Middlebrook 7H9 (BD Biosciences, 271310) medium supplemented with 0.5% v/v glycerol (Fisher Scientific, G33-4), 0.05% v/v Tween-80 (Fisher Scientific, BP338-500), and 10% v/v Middlebrook OADC (oleic acid, albumin, dextrose, catalase) (BBL, 212351).

### Aerosol infection

Equal numbers of littermates female and male *Snx5*^−/−^ and wild-type mice were infected via aerosol with a low-dose inoculum of Mtb, as previously described [47–51]. Briefly, mid-log-phase Mtb was washed three times with PBS and sonicated to disperse clumps. Then, the bacteria were resuspended in PBS (O.D._600_ of 0.1). This suspension was introduced to the nebulizer of a GlassCol aerosol chamber, calibrated to infect mice with approximately 200 bacteria per mouse. To determine the initial inoculum, on the day of infection (day zero), the entire lung from 6 mice (3 WT and 3 *Snx5*^−/−^) was collected, homogenized, and plated on 7H11 agar plates supplemented with 10% OADC (BBL, 212351), 0.5% v/v glycerol (Fisher Scientific, G30-4), and 50 µg/mL cycloheximide (Thermo Scientific, 357420050). At various time points post-infection (0, 7, 21, 42, 90, 120, and 150 days), the left lung, half of the spleen, and the left lobe of the liver were collected, homogenized, and plated on 7H11 plates for quantitation of CFU. For cytokine detection, the remaining homogenate was centrifuged and filtered. IFN-γ (Invitrogen, 88-7314-88), TNF-α (R&D Systems, MTA00B-1), IL12p40 (R&D Systems, MP400), and IL-2 (Invitrogen, 88-7024-88) were measured by ELISA according to the manufacturer’s instructions. For histology and immunohistochemistry (IHC) analysis, the superior right lung lobe was inflated and fixed in 10% buffered formalin. For survival studies, infected mice were euthanized upon reaching a loss of 15% of their maximum body weight [47–51].

### Histopathology

Lungs from infected mice were collected, fixed with 10% buffered formalin, transferred to 70% ethanol, and paraffin embedded. Then, paraffin-embedded lungs were sectioned and stained with hematoxylin and eosin (H&E) by the UT Southwestern Medical Center Histo-Pathology core facility. The slides containing lung sections were scanned using a NanoZoomer S60 digital scanner. Next, the inflammation area in each lung was quantified in a blinded analysis using ImageJ software (NIH).

### Flow cytometric assessment of lung cell population

Lungs from WT and *Snx5*^−/−^ mice uninfected or infected with Mtb were minced and incubated in PBS with 0.5% BSA, 0.2 mg/mL DNase I (Sigma, DN25), and 5mg/mL collagenase type IV (Gibco, 17102-015) on an orbital shaker at 100 rpm at 37 °C for 30 min. After digestion, the lung tissues were processed through a 70 μm cell strainer on a 50 mL conical tube. Then, the filters were washed with BSA buffer (PBS + 0.5% BSA), and the single-cell suspension was centrifuged at 350 × g for 8 min at 4 °C, and then resuspended in 1 mL of ACK lysis buffer (Gibco, A10492-01). After 1 min, BSA buffer was added to stop the reaction, and cells were centrifuged at 350 × g for 7 min at 4 °C. Cells were then transferred to a 96-well plate. After centrifugation, cells were resuspended in 50 μL of Fc-blocking solution (Invitrogen, 14-0161, 1:100) for 20 min at 4 °C. Cells were washed with staining buffer (PBS + 2.5% FBS) twice and centrifuged at 350 × g for 5 min at 4 °C. Finally, samples were incubated with surface antibody cocktail resuspended in brilliant stain buffer (Invitrogen, 00-4409-75) for 40 min at 4 °C and fixed with 1% paraformaldehyde PFA overnight. Samples were acquired on a BD LSR II at the UT Southwestern Flow Cytometry Facility. The compensation matrix was calculated using UltraComp ebeads (Invitrogen, 01-3333-42) for the antibodies and tetramers and ArC™ Amine Reactive Compensation Bead Kit (Invitrogen, A10628) for the LIVE/DEAD™ Fixable dead cell stain kits. Data analysis was performed using FlowJo 10 software. The antibody cocktail included CD4 (Super Bright 600, Invitrogen, 63-0042-82, 0.625 μL per test), CD45 (eFluor 450, Invitrogen, 48-0451-82, 2.5 μL per test), CD3 (APC-eFluor 480, Invitrogen, 47-0032-82, 5 μL per test), CD11b ( Super Bright 780, Invitrogen, 78-0112-82, 0.15 μL per test), Ly-6G (Super Bright 702, Invitrogen, 67-9668-82, Invitrogen, 0.15 μL per test), F4/80 (Alexa Fluor 488, Invitrogen, 1 μL per test, CD8 ( PE-Cyanine 7, Invitrogen, 25-0081-82, 0.15 μL per test), CD11c (PE-Cyanine 5, Invitrogen, 0.3 μL per test), Ly-6C (Alexa Fluor 700, BioLegend, 128024, 0.5 μL per test) and Live/dead fixable Aqua Blue (Invitrogen, L34957, 1:500)).

### T Cell functionality

Spleen and digested lungs from WT and *Snx5*^−/−^ mice uninfected or infected with Mtb were processed through a 70 μm cell strainer on a 50 mL conical tube and washed twice. Cells were resuspended in 1mL of RPMI with 0.5 mg/mL of anti-CD28 (BioLegend, 102102) and Golgi Plug (1:1000, BD Biosciences, 51-230KZ) and transferred to a 6-well plate, pre-coated with 1 μg/mL of anti-CD3 (BioLegend, 100302) for 5 h at 37 °C. After 5h of incubation, the cells were collected and washed 2 times with PBS containing 1% FBS. Cells were incubated in 1 mL of ACK lysis buffer for 1 min at room temperature and washed 2 times with PBS + 1% FBS. After centrifugation, cells were resuspended in 50 μL of Fc-blocking solution (Invitrogen, 14-0161, 1:100) for 15 min at 4 °C. Cells were washed and incubated with surface antibody cocktail (CD4 (Super Bright 600, Invitrogen, 63-0042-82, 0.625 μL per test), CD45 (eFluor 450, Invitrogen, 48-0451-82, 2.5 μL per test), CD3 (APC-eFluor 480, Invitrogen, 47-0032-82, 5 μL per test), and Live/dead fixable Aqua Blue (Invitrogen, L34957, 1:500)) resuspended in brilliant stain buffer (Invitrogen, 00-4409-75) for 30 min at 4 °C. Cells were then washed twice and permeabilized and fixed with BD Cytofix/Cytoperm (BD Biosciences, 554722) according to the manufacturer’s instructions. Cells were washed twice with BD Perm/ Wash Buffer (BD Biosciences, 554723) and incubated with intracellular antibody cocktail (IFN-γ (Alexa Fluor 488, Invitrogen, 53-7311-82, 1 μL per test), TNF-α (Brilliant Violet 786, Invitrogen, 417-7321-82, 1.25 μL per test), and IL-2 (PE-Cyanine 7, Invitrogen, 25-7021-82, 5 μL per test)) diluted in BD Perm/Wash Buffer for 30 min at 4 °C. fixed with 1% PFA overnight at 4 °C. Samples were acquired and data analysis were performed as described above in **Flow cytometric assessment of lung cell population.** After all washed steps, cells were centrifuged at 350 × g for 5 min at 4 °C.

### Cells and cells culture

Bone marrow cells from *Snx5*^−/−^ or wild-type mice femurs and tibia were flushed and used to generate bone marrow-derived macrophages (BMDMs) and bone marrow dendritic cells (BMDCs). To obtain BMDMs, bone marrow cells were cultivated in DMEM (Gibco, 11965-092) containing 20% heat-inactivated endotoxin-free FBS (Gibco, 1600-044), 30% L929 cell-conditioned media, Glutamax (Gibco, 35050-061), 1 mM sodium pyruvate (Gibco, 11360-070), 1 mM HEPES (Cytiva, SH30237.01), and 50 mM b-mercaptoethanol (Gibco, 21985-023). Media was replaced after 3 days of culture. After 7 days, adherent BMDMs were detached from Petri dishes (100×15mm) with 1 mM EDTA in PBS and cultured in DMEM supplemented with 10% FBS and 5% L929 cell-conditioned media. To obtain BMDCs, bone marrow cells were cultured in RPMI medium (Gibco, 11875-093) supplemented with 10% FBS (Gibco), Glutamax (Gibco, 35050-061), 1mM HEPES (Cytiva, SH30237.01), and 20 ng/mL mouse recombinant GM-CSF (Peprotech, 315-03). After 3 days of culture, additional media containing rGM-CSF was added, and after 6 days, 50% of the supernatant was replaced by fresh media containing rGM-CSF. Unattached cells were harvested on day 8 and used for experiments. RAW 264.7 murine macrophage cell line was cultured in DMEM supplemented with 10% FBS and 1mM HEPES. THP-1, a human leukemia monocytic cell line, was cultivated using RPMI 1640 media supplemented with 10% FBS, glutamax, 1 mM HEPES, and 1 mM sodium pyruvate. Cell viability was determined with a trypan-blue exclusion test, and cells were incubated overnight at 37 °C in a 5% CO_2_ incubator before use in experiments.

### Lentiviral transduction for knockdown

For *Snx5* knockdown in RAW 264.7 cells, a shRNA specific for *Snx5* (Sigma, Mission siRNA, TRCN0000381718 – GTACCGGTGCAGAGCTGCATCGACTTATCTCGAGATAAGTCGATGCAGCTCTGCA TTTTTTG) and a control scrambled sequence were obtained from Sigma. To generate lentivirus particles, HEK293T cells were co-transfected with the lentiviral vectors and psPAX2 and pMD2 (packing plasmids) at a mass ratio of 4:3:1. After 72 h, HEK293T cell supernatant was collected and purified by centrifugation at 500 × g for 5 min and then filtered using a 0.45-μm filter. Then, RAW 264.7 cells were incubated with the virus-containing supernatant and 8 μg/mL polybrene (Milipore, TR-1003-G) for 48 h, followed by puromycin selection (3 μg/mL) (Fisher Scientific, BP2956-100).

### Western blot

Cells were collected in MPER buffer (Thermo Scientific, 78501) supplemented with Protease Inhibitor (Roche, 11836170001) and incubated for 30 min at 4 °C. Next, cellular extracts were quantified using BCA assay (Thermo Scientific, 23225). 60 μg of protein from each extract was loaded onto 4-20% SDS-PAGE for electrophoresis. Proteins were then transferred to a nitrocellulose membrane using a Bio-Rad Trans-blot Turbo apparatus for 7 min at 25 V, 1.3A. The membrane was then blocked with 5% milk powder (Lab Scientific bioKEMIX, 978-907-4243) in TBST (1x TBS + 0.1% Tween 20) for 1 h at room temperature. Membranes were then incubated overnight at 4 ^◦^C with anti-SNX5 (1:400, sc-515215, Santa Cruz), anti-SNX6 (1:400, sc-365965, Santa Cruz), SNX32 (1:1000, 25763-1-AP, Proteintech) or anti-actin (1:5000, sc-47778, Santa Cruz) diluted in TBST + 1% milk solution. Each membrane was subsequently incubated with HRP-coupled anti-rabbit (Jackson, 111-035-003) or anti-mouse (Jackson, 115-035-174) IgG secondary antibodies. Blots were then developed with an ECL Chemiluminescence Kit (BioRad, 170-5060) and detected using the ChemiDoc MP Imaging System. Densitometry quantification of the bands was performed using Image J software.

### Macrophage infection

WT and *Snx5*^−/−^ BMDMs were seeded and, after 3 h, where indicated, primed with 100 UI/mL IFN-γ for 24 h. For Mtb infection, the mid-log-phase bacterial culture was washed three times with PBS. The bacterial suspension was centrifuged at 300 x g to remove bacterial aggregates and then, to generate a single-cell suspension, sonicated using Branson digital sonifier (3 times for 7s and 90% of amplitude). Cells were inoculated with Mtb, centrifuged at 1500 rpm for 10 min, and then incubated at 37°C, 5% CO_2,_ and 95% humidity. After 30 min, cells were washed twice with PBS to remove non-internalized bacteria. Fresh media with or without IFN-γ was added, and the cells were incubated at 37°C for different time points. Cell density and MOI are described in the following sections.

### Measurement of bacterial growth in macrophages

WT and *Snx5*^−/−^ BMDMs were seeded in 48-well plates (2×10^5^), with or without IFN-γ (100 UI/mL) priming (R&D Systems, 485-MI), and infected with Mtb (MOI of 1). Cells were lysed on the same day of infection (day 0) and 1, 3, and 5 days post-infection with 0.5% Triton X-100 (Sigma, T8787) in water. Serial dilutions were plated on 7H11 plates, and colony-forming units (CFUs) were counted after 3 weeks of incubation.

### Binding and phagocytosis assay

The effect of *Snx5* knockout on the binding and phagocytosis of Mtb was evaluated by CFU and by microscopic quantification. For the CFU experiment, BMDMs were plated in 48-well plates (2×10^5^), while for microscopy, cells were seeded in 24-well plates (2×10^5^) containing coverslips. Initially, the cells, with or without IFN-γ (100 UI/mL) priming, were incubated for 10 min at 4 °C. Then, they were incubated with mCherry-Mtb (MOI of 1), centrifuged at 4 °C for 10 min at 500 x g, and incubated at 4°C for another 30 min; this step allows binding without bacterial phagocytosis. After this period, the cells were washed to remove non-internalized Mtb. For the binding study, the cells were fixed with1% PFA overnight at 4°C or lysed with 0.5% Triton X-100, as described in the ‘**Measurement of bacterial growth in macrophages’**. For the phagocytosis study, the cells were re-incubated with DMEM at 37°C. After 1 h, the cells were washed and fixed with 1% PFA overnight at 4°C or lysed with 0.5% Triton X-100. Fixed cells were labeled with 488-wheat germ agglutinin (WGA) (1:1000) (Invitrogen, W11261) for 1 h, then mounted using ProLong Diamond Antifade kit containing DAPI (Invitrogen, P36962), protected from light, and cured for 24 h at room temperature. Then, images were acquired on a Zeiss AxioImager Z2. For quantification, triplicate samples were analyzed for each condition, and a minimum of 100 cells were counted per sample using Imaris (7.4.0 version; Bitplane).

### Immunofluorescence

WT and *Snx5*^−/−^ BMDMs were plated in 24-well plates (2×10^5^) containing coverslips, with or without IFN-γ (100 UI/mL) priming, and infected with mCherry-Mtb (MOI of 1). After 24 h, infected cells were fixed with 1% PFA overnight at 4 °C, and then transferred to PBS. Cells were permeabilized with cold methanol for 20 min at –20 °C, washed three times in PBS, and blocked with 3% BSA for 1 h at room temperature. Next, the cells were stained with primary antibodies in 3% BSA overnight at 4°C. The primary antibodies used were anti-EEA1 (Abcam, ab2900, 1:250), and anti-LAMP (Novus Biologicals, NB120-19294, 1:50). Alexa-Fluor 488 or 647-conjugated donkey anti-rabbit (1:1000 or 1:500, respectively) was used as a secondary antibody. Finally, the coverslips were mounted using ProLong Gold Antifade kit containing DAPI (Invitrogen, P36962), protected from light, and cured for 24 h at room temperature. Z-stack images were acquired with a Nikon CSU-W1 SoRa spinning disk confocal microscope using a 100×/1.4 objective. Triplicate samples were quantified for each condition in a blinded analysis, and a minimum of 100 bacteria were counted per sample using Imaris (7.4.0 version; Bitplane).

### Cytokine quantification

To assess cytokine production in Mtb-infected cells, WT and *Snx5*^−/−^ BMDMs (10^6^) were seeded in 12-well plates (10^6^), primed or not with IFN-γ (100 UI/mL), and infected with Mtb (MOI of 5). After 24 h, the supernatant was collected and filtered (Pall Corporation, 8582). MCP-1 (Biorad, 171G5019M), IL-6 (R&D Systems, M6000b), TNF-α (R&D Systems, MTA00B), and IL-12p40 (R&D Systems, MP400) were measured by ELISA according to the manufacturer’s instructions.

### Endolysosomal proteolytic assay

Proteolytic activity in BMDMs was evaluated using DQ-BSA (DQ^TM^ Green BSA, 2540586), a modified bovine serum albumin that fluoresces upon cleavage in hydrolytic compartments, such as lysosomes. WT and S*nx*5^−/−^ BMDM were plated in 24-well plates (2×10^5^) containing coverslips, primed or not with IFN-γ (100 UI/mL), and infected with mCherry-Mtb (MOI of 1). After 24 h, cells were incubated with DQ-BSA (10 μg/mL) for 30 min. The cells were fixed with 1% PFA overnight at 4 °C, and then transferred to PBS. Coverslips were mounted, and images were acquired as described above in **Immunofluorescence**. The intensity of DQ-BSA per pixel for at least 50 cells was analyzed using Imaris (7.4.0 version; Bitplane).

### Antigen presentation assay

BMDMs, DCs, or RAW 264.7 cells were plated in 96-well (5×10^4^) round-bottom plates (Thermo Scientific, 163320) with or without IFN-γ (100 UI/mL) priming. After overnight incubation, the cells were fed with vehicle or OVA_323-339_ (the epitope of ovalbumin (OVA) presented by I-Ab, 5 mg/mL) (InvivoGen, 5825-43-01), Ag85B (10ng/mL) (Ray-Biotech, 228-11030 (discontinued) or BEI, NR5326), or heat-killed Mtb (MOI of 10) (Table 1). Then, cells were washed with PBS and co-cultured with antigen-specific MHC class II-restricted T cell hybridomas (10^5^) (Table 1) in a medium supplemented with 1 mM aminoguanidine hemisulfate (BioSynth, FA170452 and Santa Cruz, sc-202930). After co-incubation, the supernatants were collected, and T cell activation was measured using IL-2 ELISA (Invitrogen, 88-7024-88).

**Table 1.**
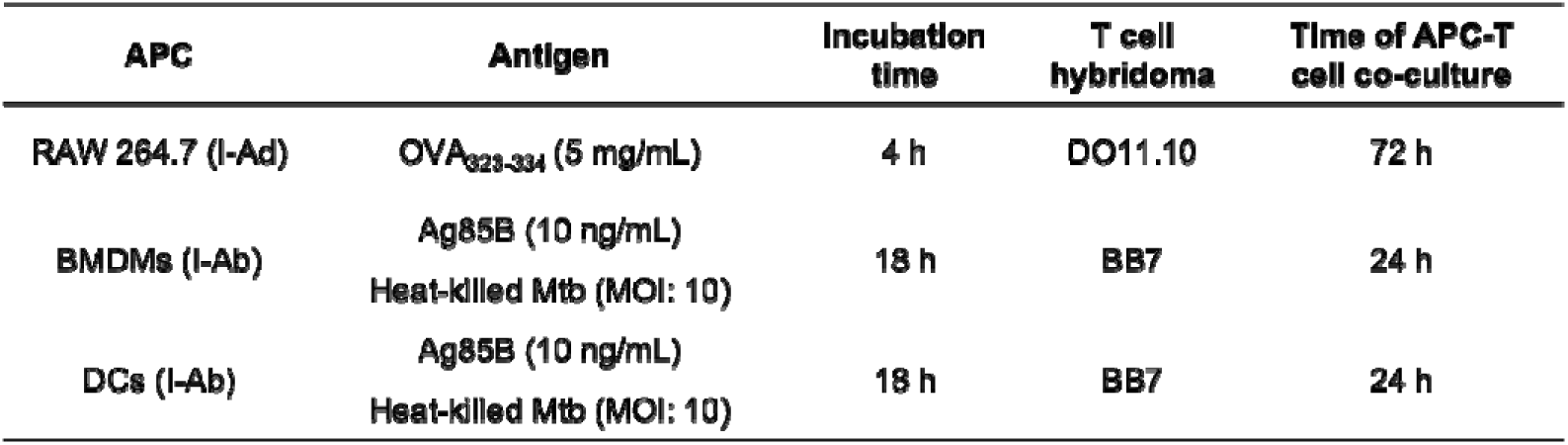
Antigen presentation assay experimental set-up.

### Flow cytometry

BMDMs (10^6^), primed or not with IFN-γ (100 UI/mL), were washed twice with PBS and incubated with 1% BSA at 4 °C for 30 min to reduce non-specific binding. Then, the cells were labeled with surface markers specific to MHC-II (APC, R&D Systems, FAB6118A, 10 μL per test), CD80 (FITC, BD Biosciences, 553768, 0.25 μg per test), or CD86 (PE, BD Biosciences, 553692, 0.06 μg per test) at 4°C for 30 min. Subsequently, the cells were washed again with PBS and fixed with 1%PFA. 20,000 events were acquired on the BD FACSCalibur. Data analysis was performed using FlowJo 10 software.

### Click chemistry

To compare the amount of peptide conjugated to MHC-II on the cell surface of BMDMs, we adapted a published protocol [52]. Of note, this protocol uses an extended influenza A virus hemagglutinin (HA) peptide (HA_318-338_), which must be cleaved and loaded onto MHC-II for presentation on the cell surface. This peptide was designed based on the crystal structure of the HA_322-334_ association with the human HLA-DR1 MHC class II. Due to technical limitations in using PMA-activated THP-1 cells (human macrophage-like cells), we used BMDMs in our experiments. WT or *Snx5*^−/−^ cells (10^6^) were plated in 35×10 mm Petri dishes, primed with IFN-γ (100 UI/mL) for 24 h, and incubated with 50 mM of influenza HA_318-338_ in FBS-free media. Control cells were incubated with non-modified peptide (YGACPKYV**K**QNTLKLATGMRN – non-clickable), and testing cells were incubated with a clickable version of the HA peptide, in which the lysine (K326) was converted into a propargylglycine (pra), a non-naturalized amino acid compatible with bio-orthogonal labeling using CalFluor488 (YGACPKYV**(pra)**QNTLKLATGMRN – clickable) (synthesized by GenScript). After 18 h, cells were gently detached by flushing with PBS and stained with Live/dead fixable Aqua Blue (Invitrogen, L34957, 1:500) to assess cell viability. Cells were then fixed with 4% PFA for 15 min and incubated with quenching buffer (100 mM glycine and 100 mM NH_4_Cl in PBS) for 20 min to reduce autofluorescence. Next, the surface presentation of HA was labeled with 100 mM of CalFluor488 (Broadpharm, BP-24164) (diluted in a buffer containing 1 mM CuSO4, 0.5 mM THPTA (Tris(3-hydroxypropyltriazolylmethyl)amine), 5 mM sodium L-ascorbate, 0.01% BSA, and aminoguanidine 2.2%) for 1 h. Surface MHC-II were subsequently labeled with APC-conjugated anti-MHC-II (R&D Systems, FAB6118A, 10 μL per test). Samples were acquired on a BD FACSCanto, and data analysis was performed using FlowJo 10 software.

### RNA isolation for RNA sequencing

WT and *Snx5*^−/−^ BMDMs from 3 different animals were plated in 6-well plates (10^6^) and infected or not with Mtb (MOI of 5). After 4 or 24 h of infection, samples were lysed using TRIzol (Invitrogen, 15596018) for 5 min, incubated with chloroform, and centrifuged at 12,000 x g for 5 min at 4 °C. Then, the aqueous phase was subject to RNA extraction using the RNeasy Mini Kit (QIAGEN, 74104) according to the manufacturer’s protocol.

### RNA sequencing data acquisition and analysis

RNA samples were transferred to McDermott Center Next Generation Sequencing (NGS) Core at UT Southwestern Medical Center for RNA quality control evaluation, cDNA library preparation, and sequencing. The RNA quality was confirmed using Agilent Tapestation 4200 (RNA integrity number ≥ 8.0), and the initial RNA concentration was determined using Qubit. A total of 1 mg of DNAse-treated RNA was used for library preparation using Illumina’s TruSeq Stranded mRNA Library Prep Kit. Sequencing was performed using Illumina NextSeq 2000 with a sequencing depth of 25-35M reads and 75 bp single-end reads. Then, RNA-Seq reads were aligned to the *Mus musculus* reference genome. Finally, downstream analyses were performed using iDEP (integrated Differential Expression and Pathway analysis) online tools [53]. Briefly, the data were preprocessed to remove low level expressed genes, and differential expression analysis between groups was performed using the DESeq2 R package with a false discovery rate (FDR) threshold of 0.1. Genes with a fold change > 2 found by DESeq2 were considered significant.

### Statistical Analysis

Statistical analysis and graphing were performed using GraphPad Prism software (version 10). After verifying data normality, the unpaired Student’s t-test, or the Mann-Whitney test was used to compare two groups. Comparisons among three or more groups were assessed with one-way ANOVA for Gaussian distributions or the Kruskal-Wallis nonparametric test. Differences among groups were considered statistically significant when *p* < 0.05.

## Acknowledgments

The authors would like to thank Xiaonan Dong, Cody Ruhl and Haaris Khan for preliminary experiments with *Snx5^−/−^* mice. We are grateful to Dr. Sung-Moo Park for assistance with antigen presentation assays and to Dr. Christina Stallings for flow cytometry guidance. Many thanks to the UT Southwestern Quantitative Light Microscopy Core, particularly the core director Marcel Mettlen. We thank the UTSW Genomics Sequencing Core for bulk RNA-seq library preparation and sequencing. The Histopathology Core and the CRI Flow Cytometry Facility, especially senior specialist Fernanda Lins, are acknowledged for their technical assistance. We also thank Dr. Denise Ramirez and the Whole Brain Microscopy Facility at UT Southwestern. We thank BEI Resources, NIAID, NIH, for providing Ag85B used in this study. NIH 1S10OD028630-01 to Dr. Kate Luby-Phelps was used to purchase the Nikon CSU-W1 with SoRa (spinning disk confocal microscope) used in this work. We acknowledge the Imaris customer support team for their guidance in developing a pipeline for fluorescence intensity quantification.

## Competing interest

All authors declare that they have no competing interests.

## Funding

This work was supported by the National Institutes of Health U19 AI142784 (M.U.S), T32HL098040 (K.C.R.) and T32 AI007520 (K.F.N).

## Data availability

All RNA-Seq data have been deposited in the Gene Expression Omnibus (GEO) under accession number: GSE317798.

## Author’s contributions

Conceptualization, B.R.S.D., M.U.S.; Formal analysis, B.R.S.D., M.U.S.; Sample collection, B.R.S.D., K.F.N., V.A.E.; Investigation, B.R.S.D., K.F.N., V.A.E., S.A.A, K.C.R, P.C.C; Funding acquisition, K.F.N., K.C.R., M.U.S.; Project administration, M.U.S.; Supervision, M.U.S.; Writing – original draft, B.R.S.D., M.U.S.; Writing – review and editing, All authors.

## Notes

### Competing Interest Statement

The authors have declared no competing interest.

