## Supplemental Figures and Legends for "Sorting nexin 5 mediates antigen presentation and immunity against *Mycobacterium tuberculosis*"

Supplemental Fig. 1

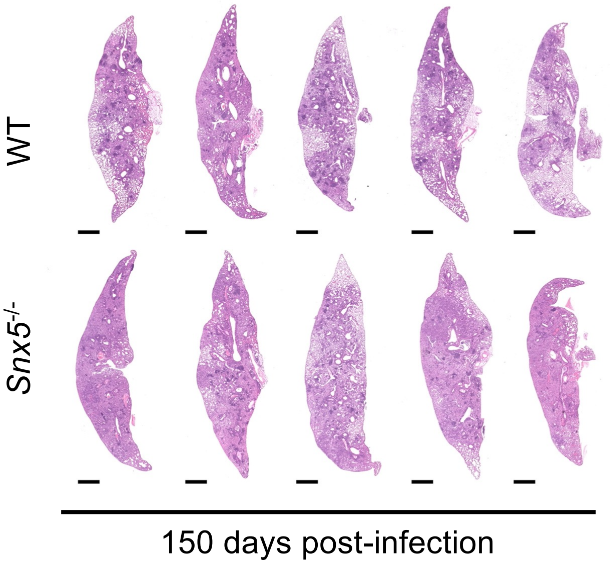

### Supplemental Fig. 2

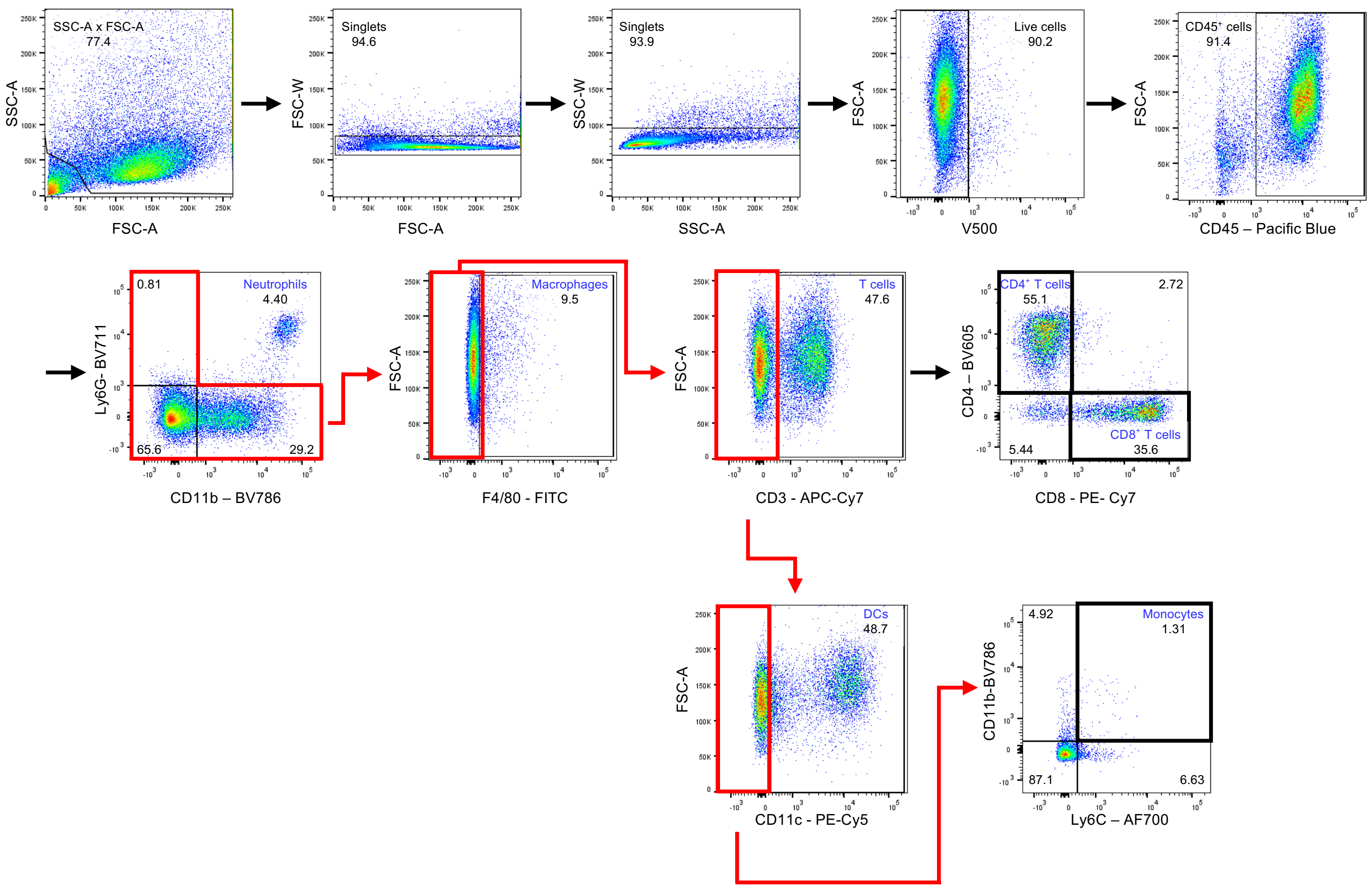

Supplemental Fig. 3

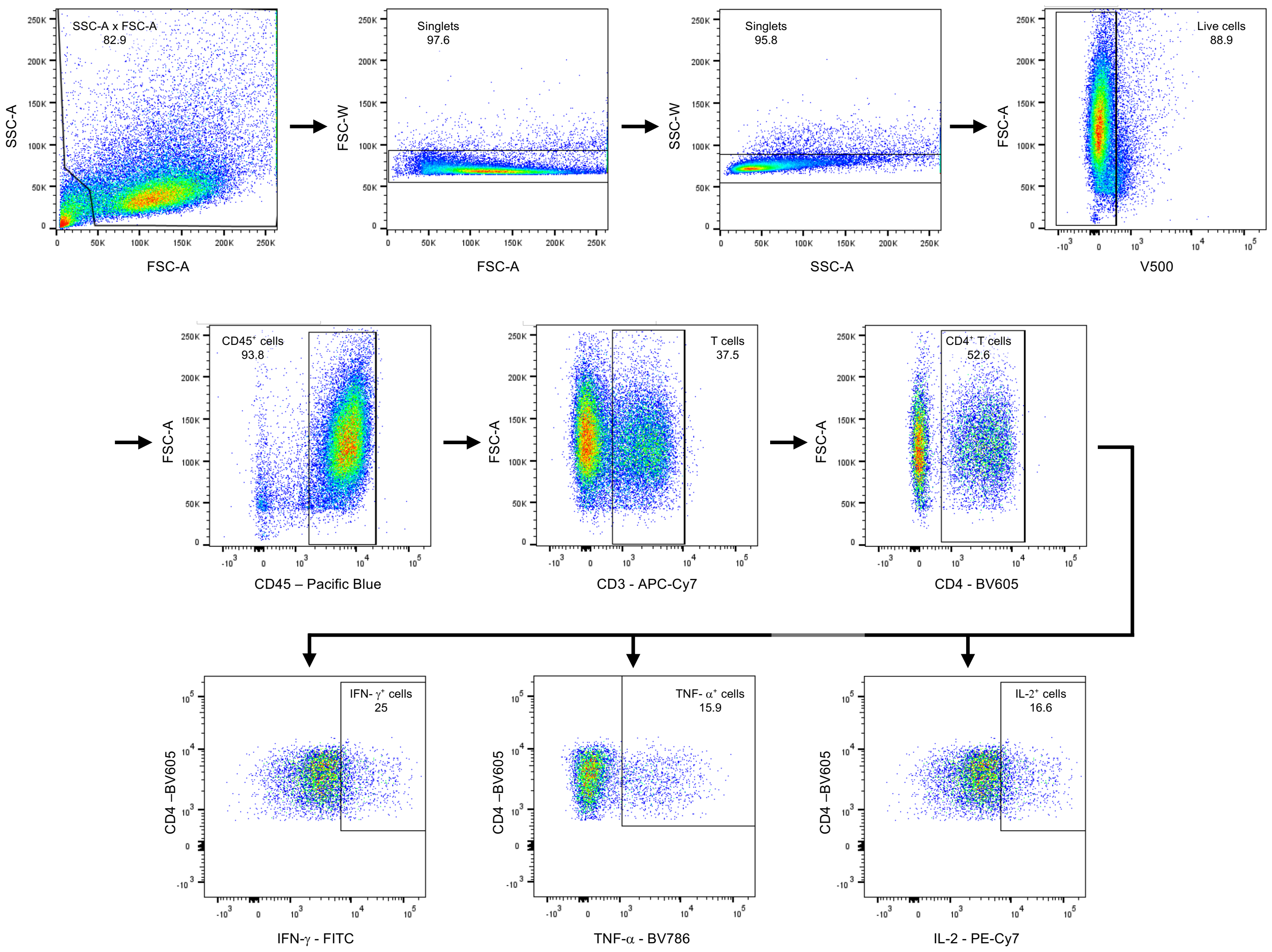

Supplemental Fig. 4

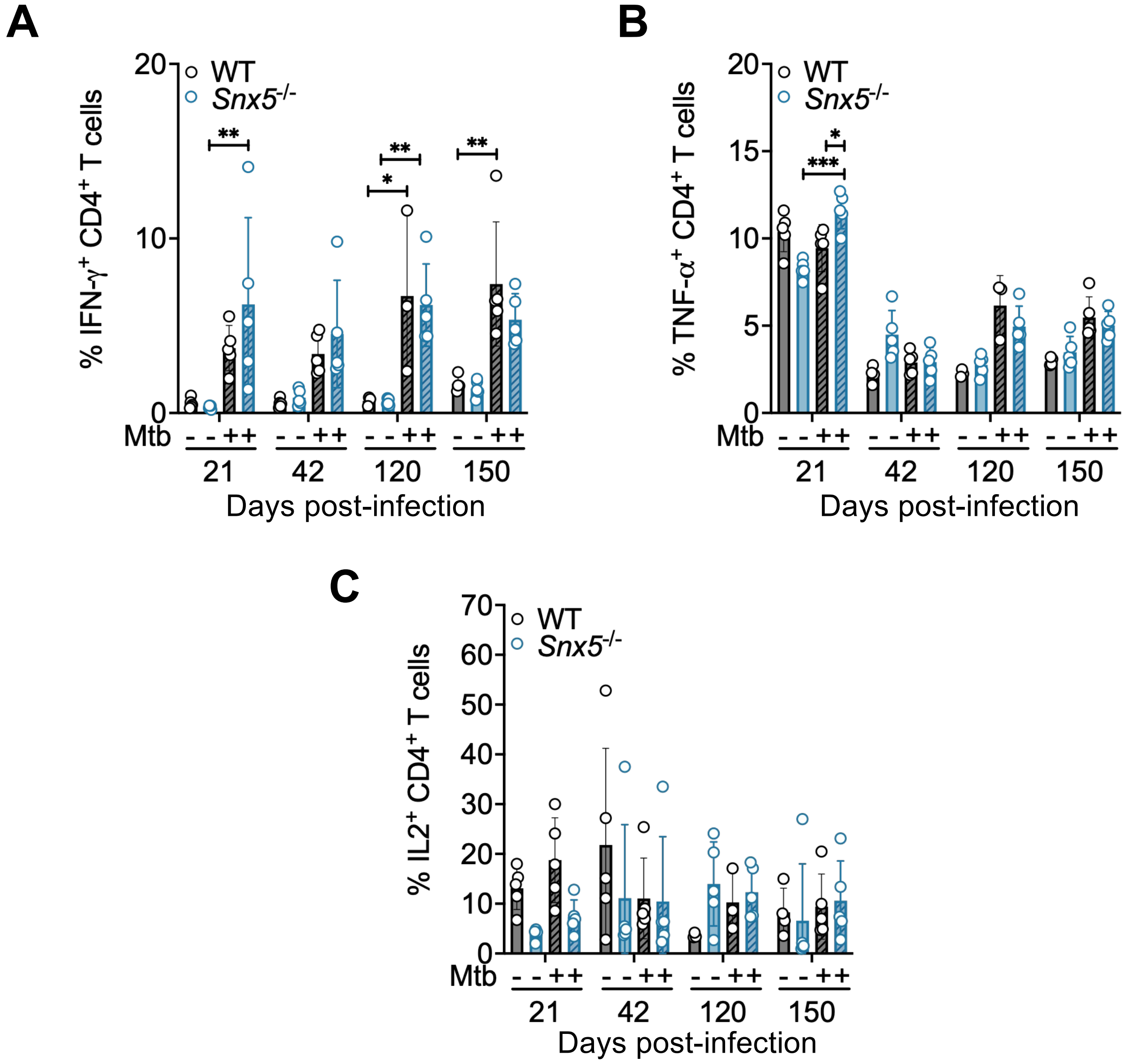

Supplemental Fig. 5

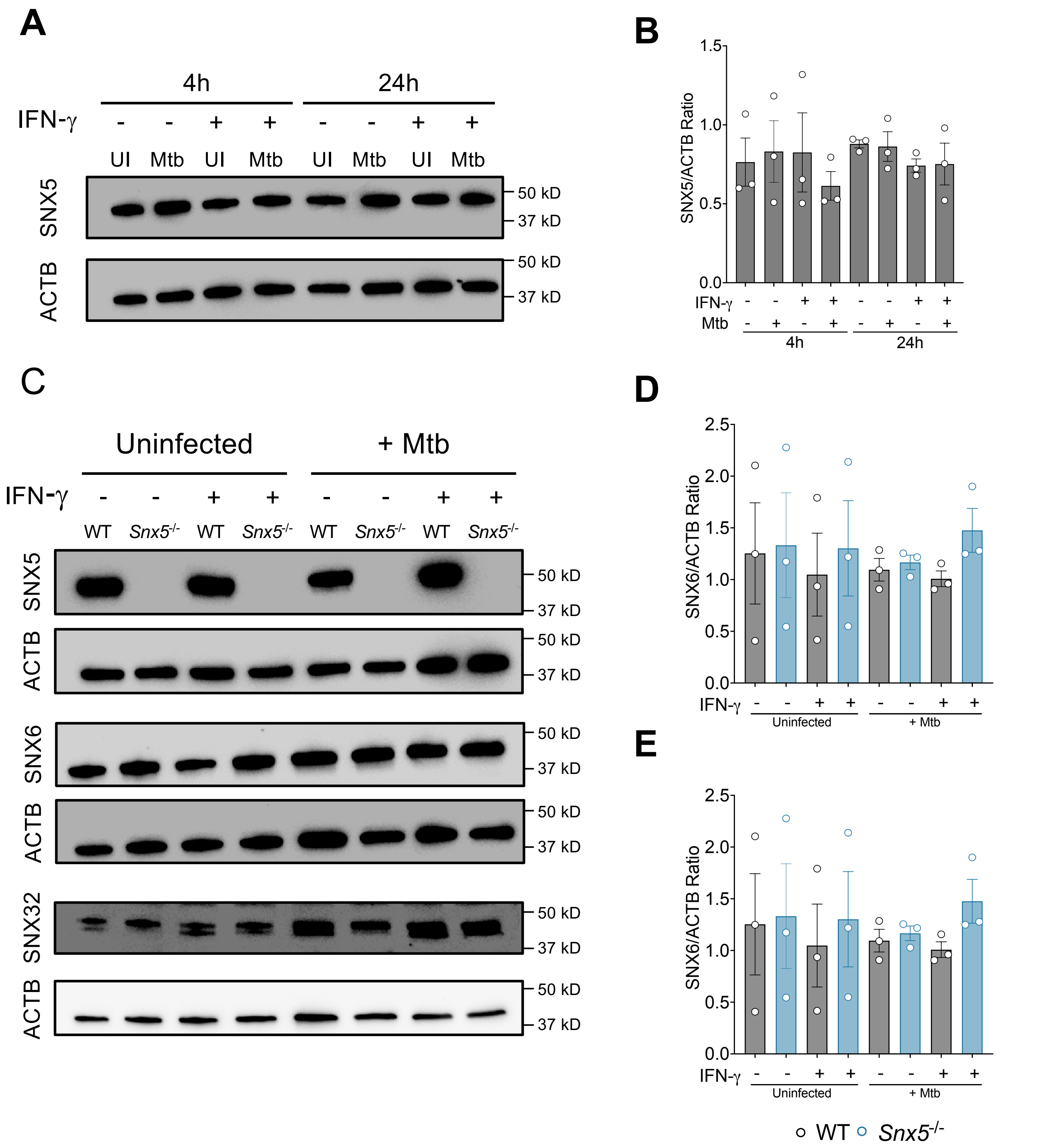

### Supplemental Fig. 6

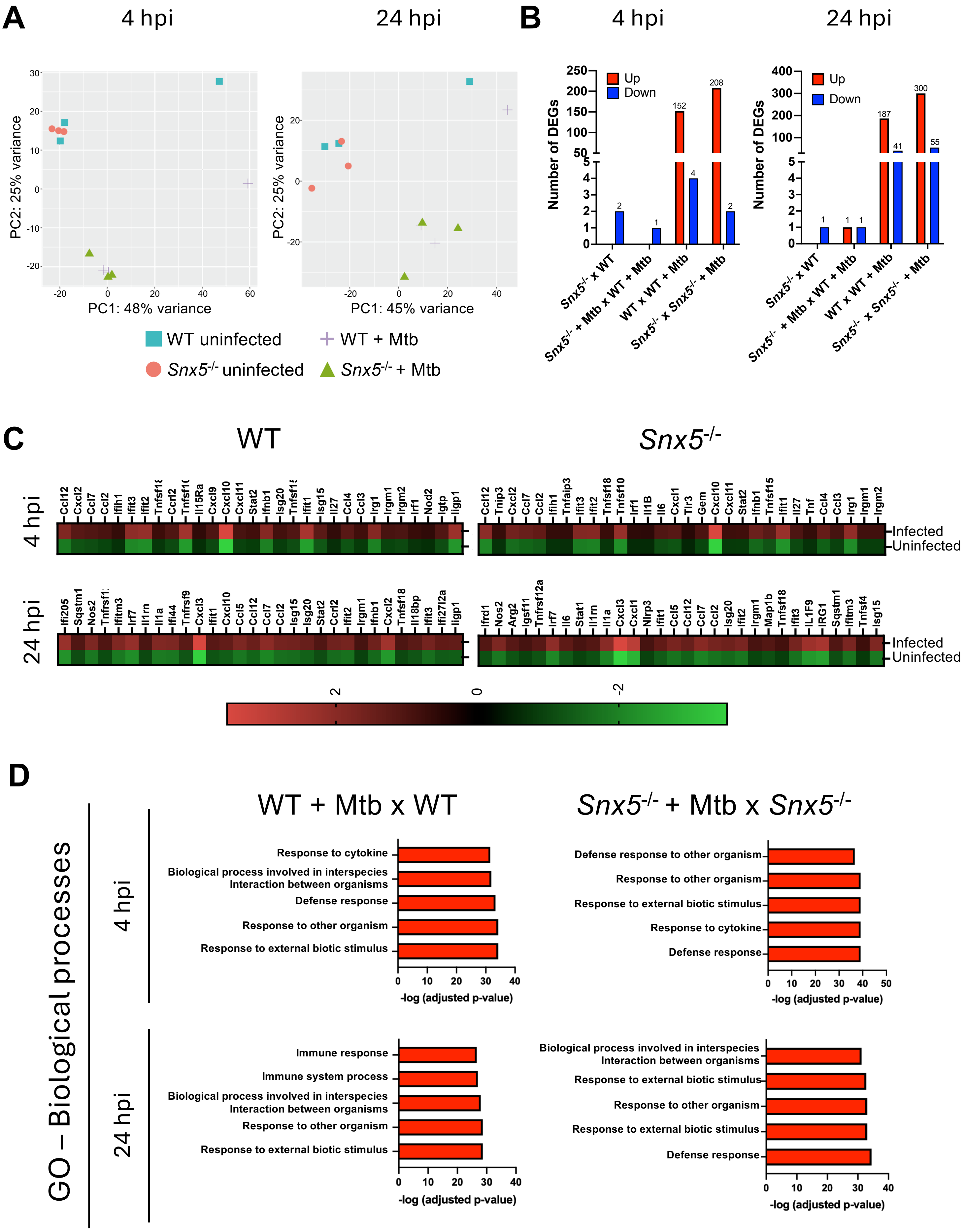

Supplemental Fig. 7

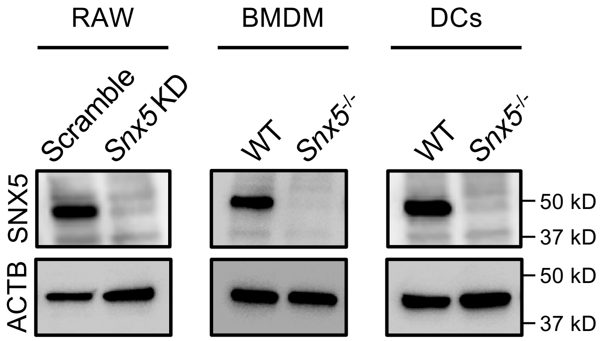

### Supplemental Fig. 8

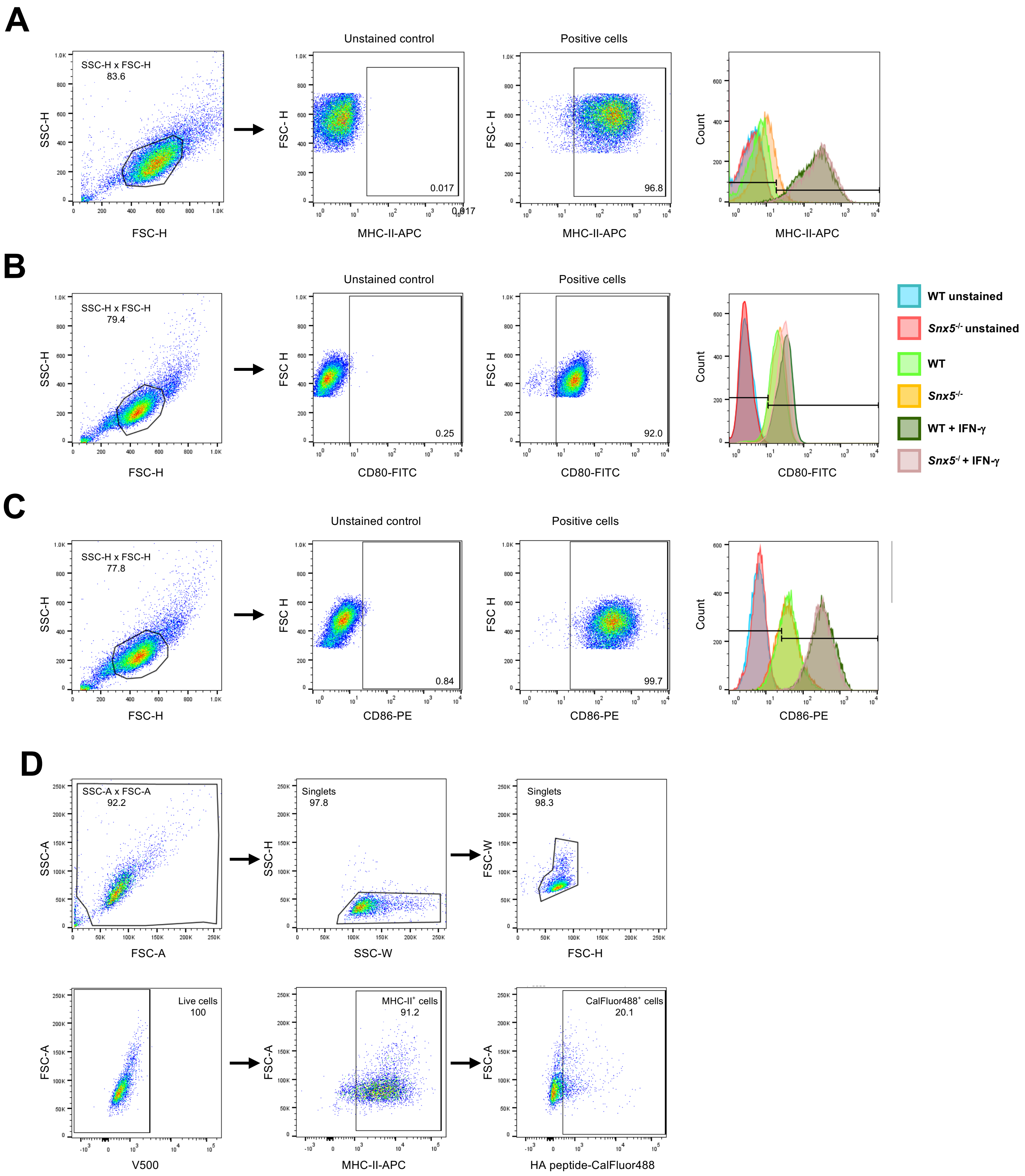

**Supplemental Figure 1 related to Figure 2: Comparison of Mtb-induced pulmonary inflammation in WT and *Snx5*$^{-/-}$ mice.** Photomicrographs of H&E-stained lungs at 150 days post-infection with Mtb. Scale bars, 1000 μm.

**Supplemental Figure 2 related to Figure 2: Gating strategy for flow cytometric analysis of percentage of immune cells in the lung of Mtb-infected mice.** Isolated leukocytes from total lung of WT and *Snx5*$^{-/-}$ mice were first gated based on forward and side scatter properties, followed by selection of singlets and viable cells, cells negatively stained with Aqua Blue fixable viability dye (V500). Cells were then hierarchically sub-gating strategy to determine the percentage of neutrophils (Ly6G$^{+}$ CD11b$^{+}$), macrophages (F4/80$^{+}$), T cells (CD3$^{+}$), CD4$^{+}$ T cells (CD3$^{+}$ CD4$^{+}$), CD8$^{+}$ T cells (CD3$^{+}$ CD8$^{+}$), DCs (CD11c$^{+}$), and monocytes (CD11b$^{+}$ Ly6C$^{+}$). Black arrows indicate sequential gating and red arrows indicate Boolean NOT gates.

**Supplemental Figure 3 related to Figure 3: Gating strategy for flow cytometric analysis of the production of pro-inflammatory cytokines by CD4$^{+}$ T cells.** Isolated leukocytes from the total lung of WT and *Snx5*$^{-/-}$ mice were hierarchically selected based on forward and side scatter properties, followed by selection of singlets and viable cells, which were negatively stained with Aqua Blue fixable viability dye (V500). Then, CD45-, CD3-, and CD4-positive cells were selected. CD4$^{+}$ cells were further gated for pro-inflammatory cytokine expression (TNF-α, IFN-γ, and IL-2).

**Supplemental Figure 4 related to Figure 3: *Snx5* deletion does not alter the production of cytokines in splenic T cells.** Explanted splenic CD4$^{+}$ T cells were characterized for cytokine expression, including (A) IFN-γ, (B) TNF-α, and (C) IL-2 using surface and intracellular cytokine staining for flow cytometry. Each circle corresponds to one animal. Bars are the mean ± S.D. for 3 to 5 animals per group per time point. *$p < 0.05$, **$p < 0.01$ and ***$p < 0.001$ (for indicated comparison; one-way ANOVA).

**Supplemental Figure 5 related to Figure 4: Detection of SNX5, SNX6, and SNX32 in Mtb-infected BMDM**. WT and *Snx5*$^{-/-}$ BMDMs, either primed or not with IFN-γ, were infected with Mtb Erdman (MOI: 5). (A) Western blot of SNX5 and Actin (ACTB) expression in WT BMDMs infected or not with Mtb for 4 or 24 h. (B) Densitometric quantification of SNX5/Actin ratios. (C) Western blot of SNX5, SNX6, SNX32, and Actin expression in WT and *Snx5*$^{-/-}$ BMDMs infected or not with Mtb for 24 h. (D and E) Densitometric quantification of (D) SNX6/Actin and (E) SNX32/Actin ratios. ACTB: Actin. Symbols represent individual experiments, while lines are representative of means ± SE.

**Supplemental Figure 6 related to Figure 4: *Snx5* does not affect macrophage transcriptional responses to Mtb infection**. WT and *Snx5*$^{-/-}$ BMDMs were infected with Mtb Erdman for 4 or 24 h, and transcriptomes were analyzed by RNA sequencing (A) Principal-component analysis of either infected or not with Mtb WT or *Snx5*$^{-/-}$ BMDMs. (B) Number of significantly differentially expressed genes (DEGs) in either infected or not WT x *Snx5*$^{-/-}$, WT x Mtb-infected WT, and *Snx5*$^{-/-}$ x Mtb-infected *Snx5*$^{-/-}$ (FDR < 0.1, FC > 2). (C) Heat map comparing the $\log_2$ fold change of 30 differentially expressed genes during Mtb infection. (D) Top 5 GO Biological Processes. KO: *Snx5*$^{-/-}$; + Mtb: Mtb-infected BMDMs.

**Supplemental Figure 7 related to Figure 8: Western blot validation of SNX5 knockdown in RAW cells and knockout in BMDMs and DCs.** Representative western blot of SNX5 accumulation in RAW cells encoding a scramble or *Snx5*-targeting shRNA, and in BMDMs and DCs from wild-type (WT) and *Snx5*$^{-/-}$ mice. KD: Knockdown. ACTB: Actin.

**Supplemental Figure 8 related to Figure 9: Gating strategy for flow cytometric analysis of antigen presentation molecules on the BMDM cell surface**. (A-C) WT and *Snx5*$^{-/-}$ BMDMs were selected based on forward and side scatter properties, followed by selection of (A) MHC-II, (B) CD80, and (C) CD86 positive cells. Representative histograms show cell-surface levels of MHC-II, CD80, and CD86 in WT and *Snx5*$^{-/-}$ BMDMs, either primed or not with IFN-γ. (D) WT and *Snx5*$^{-/-}$ BMDMs were hierarchically selected based on forward and side-scatter properties, followed by singlet and viable cell selection, which were negatively stained with Aqua Blue fixable viability dye (V500). Then,

MHC-II (APC)-positive cells were selected and further gated for HA-peptide (CalFluor 488).
